# *De novo* H3.3K27M-altered Diffuse Midline Glioma in human brainstem organoids to dissect GD2 CAR T cell function

**DOI:** 10.64898/2025.12.15.694282

**Authors:** Nils Bessler, Amber K.L. Wezenaar, Hendrikus C.R. Ariese, Celina Honhoff, Ellen J. Wehrens, Noëlle Dommann, Cristian Ruiz Moreno, Thijs van den Broek, Raphaël V.U. Collot, D.J. Kloosterman, Farid Keramati, Mieke Roosen, Sam de Blank, Esmée van Vliet, Mario Barrera Román, Lucrezia C.D.E. Gatti, Ali Ertürk, Jürgen Kuball, Zsolt Sebestyén, Marcel Kool, Sara Patrizi, Evelina Miele, Annette Künkele, Mariëtte E.G. Kranendonk, Annelisa M. Cornel, Stefan Nierkens, Christian Mayer, Hendrik G. Stunnenberg, Anna Alemany, Maria Alieva, Anne C. Rios

## Abstract

Diffuse midline glioma (DMG) is a rare yet highly aggressive paediatric cancer primarily arising in the pontine region of the brainstem, necessitating the development of scalable patient-representative models for treatment advance^1,2^. Here, we developed an FGF4-driven human brainstem organoid model, with high representation of pontine glial lineages. By genetically engineering *de novo* H3.3K27M-altered DMG, we show that this brainstem glial specification is essential for driving DMG tumorigenesis, resulting in tumours that recapitulate the infiltrative nature and molecular heterogeneity of patient samples. By performing prolonged GD2 CAR T cell treatment in this model, we could mirror variable treatment outcomes as observed in the clinic^3,4^ and demonstrate a high level of CAR T cell transcriptional heterogeneity. From these CAR T cell functional states, we could identify the most potent effector population and validated NCAM1 as a selection marker for their enrichment. In contrast, NCAM1^-^ cells were linked to a cellular stress response, previously associated to immunotherapy resistance^5^. Furthermore, incorporating the brain-resident myeloid compartment resulted in DMG-specific, largely immunosuppressive microglia subtypes^6^. These disease-representative microglia reduced GD2 CAR T cell treatment efficacy and we identified the functional profiles most susceptible to this microglia-dependent immune modulation. Thus, we present a scalable human DMG model with critical applications towards understanding CAR T cell functionality to aid therapy development for this detrimental disease.

## Main

Diffuse midline gliomas (DMGs) are rare and aggressive paediatric brain tumours often caused by somatic mutations in histone 3 (H3) genes, commonly a lysine27-to-methionine (K27M) substitution^7^ and occurring at a high prevalence in the pons region of the brainstem^1^. Primarily, affecting children under 10 years old^8^, they present the highest mortality rate of any cancer, with a median overall survival of only 9-15 months^2,9,10^. This detrimental prognosis underscores a critical need to gain more insight into the unique biology of the disease to develop more effective treatments.

Efforts to unravel the cellular composition of H3K27M-altered DMG through single-cell analysis have revealed intratumoral heterogeneity, with a spectrum of tumour cell profiles ranging from stalled stem-like oligodendrocyte progenitor cell (OPC-like) to more differentiated astrocyte (AC-like) and oligodendrocyte (OC-like) phenotypes, which closely resemble normal developmental cell types, alongside a recently identified mesenchymal-like (MES-like) state^11–13^. In addition, insights from both animal studies^14–17^ and human pluripotent stem cell-derived research^18,19^ suggest an early neurodevelopmental window of tumour initiation. Thus, dysregulated mechanisms during hindbrain development^15,20,21^, particularly involving glial neural progenitors in the region responsible for brainstem pons formation^22^, likely play a central role in driving H3K27M-altered gliomagenesis. Capturing this region-specific embryonic patterning is therefore crucial for accurately modelling pontine DMG.

Human brain organoids have become valuable *in vitro* tools for investigating brain development and understanding the onset, progression, and potential therapeutical targeting of nervous system disorders, including cancer^23–29^. Given the rarity and inoperable nature of DMG, which limits the availability of patient material^1^, organoids could offer a scalable model for generating DMG tumours *de novo* and enabling *in vitro* testing of emerging therapies. This includes the latest advances in immunotherapy for DMG; GD2 Chimeric Antigen Receptor (CAR) T cells^2,30,31^ that in a recent first in-patient clinical trial showed highly promising yet variable treatment outcomes between patients^3^. Correlative data from this trial suggest that a rise in the immunosuppressive myeloid compartment coincides with unfavourable treatment outcomes^4^. Uncovering the functional profiles of CAR T cells and their interplay with the immunosuppressive tumour microenvironment could, therefore, provide critical insights for developing strategies to further advance CAR T cell treatment outcomes in DMG.

Here, we report a novel human cerebral guided organoid model for the brainstem region, enriched for pontine-medulla glial lineages. Genetic modelling of H3.3K27M-altered DMG in these Brainstem-regionalized Organoids (BrOs), unlike unguided cerebral organoids, successfully replicates the infiltrative nature and transcriptomic landscape of DMG in patients. We demonstrate the utility of this new accessible human DMG organoid model (DMGO) for modelling CAR T cell functional heterogeneity during prolonged treatment (up to 1 month) and within the context of the brain-resident immune microenvironment.

### Brainstem organoid patterning

To create a human organoid with the right identity for subsequent DMG tumour induction and treatment response modelling, we implemented morphogen guidance based on a timely sequence of Wnt, dual SMAD inhibitors, retinoic acid (RA), fibroblast growth factors (FGFs) and sonic hedgehog (SHH) to specify hindbrain identity (**Fig. 1a, Extended Data Fig. 1a, b, Supplementary Table S1a**). While FGF2 and FGF8 can be used in combination with RA and Wnt to pattern midbrain^32^, cerebellum^33^, or spinal cord^34^ in growing organoids, we evaluated FGF4 because of its role in specifying rostral hindbrain, particularly in prepontine and pontine areas^35,36^, as well as its involvement in the development of hindbrain-specific serotonergic neurons^37^. A direct comparison of replacing common FGF2 supplementation with FGF4 after 7 days of patterning, demonstrated that 10 ng/ml of FGF4 specifically promotes developing pontine, including prepontine to retropontine area based on bulk sequencing data (**Fig. 1b, Supplementary Table S2a**). Furthermore, among HOX genes important for hindbrain formation, expression of *HOXB1*, a marker of pontine precursor cells^38^, emerged already at an early stage (day 14) (**Extended Data Fig. 1c**) and 3D imaging revealed HOXB1 expressing cells within early neurodevelopmental SOX2+ neural rosette structures (**Fig. 1c**). Importantly, bulk sequencing analysis from day 7 to day 84 showed that the patterning remained consistent and reproducible across and within multiple batches, as well as between human embryonic stem cell (hESC) and induced pluripotent stem cell (iPSC) sources (**Extended Data Fig. 1d, e**).

**Figure 1.**
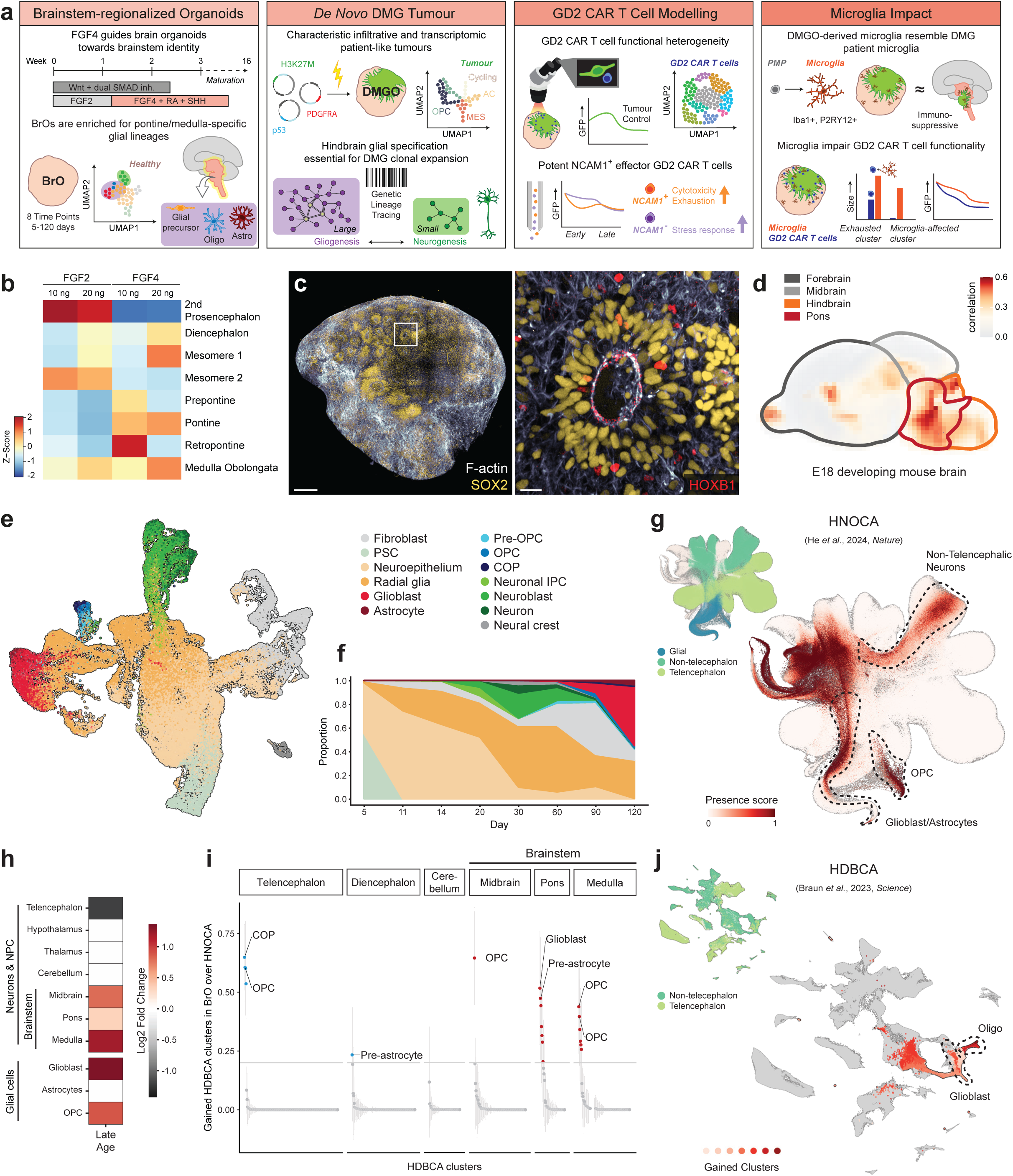
Generation and validation of glial-enriched brainstem organoids. **a**, Schematic representation of timely morphogen stimulated patterning of hESCs/hIPSCs towards brainstem organoids and their subsequent application for DMG tumour-, CAR T cell treatment-, and microglia-enriched tumour microenvironment modelling. **b**, Heatmap of z-score measuring relative brain region identity based on VoxHunt similarity mapping for various supplemented concentrations of FGF2 or FGF4. **c**, Immunofluorescent 3D images of a 200 µm thick organoid slice at day 21 labelled for F-ACTIN (white), SOX2 (yellow) and HOXB1 (red). White insert indicates zoom area displayed on the right. Overview image and zoom scale bars 250 µm and 25 µm respectively. **d**, VoxHunt spatial correlation map of day 120 brainstem organoids with E18.5 mouse brain, pons area delineated in red. **e**, Integrated UMAP representation of developing brainstem organoids from different timepoints, coloured by cell annotation. **f**, Areaplot following the relative distribution of cell types over time. Cell types colour-coded as in e. **g**, UMAP of the HNOCA^40^ coloured for brainstem organoid presence score. A high score indicates a high likelihood that these HNOCA cells are present in the brainstem organoid dataset. Areas annotated by a dashed line indicate lineages as annotated in the HNOCA. Insert UMAP is coloured by coarse regional annotation. **h**, Heatmap showing the log2 fold compositional changes in the brainstem organoid dataset compared to the HNOCA, positive values correspond to an increased abundance of cells from the indicated regional identity or glial lineage. **i**, Cell clusters in the HDBCA^41^ with gained coverage in brain stem organoids relative to the HNOCA. The horizontal line indicates the threshold used to define a cluster as gained or not. **j**, UMAP of the HDBCA with in shades of red the HDBCA clusters gained in brainstem organoids, mostly related to Oligo and Glioblasts. Grey represents clusters below the threshold used to define gained. Insert UMAP is coloured by coarse regional annotation.

To investigate cellular composition and regional identities at higher resolution, we performed time-course single-cell RNA sequencing (scRNA-seq) across eight-time points, spanning from day 5 to day 120 (**Extended Data Fig. 2a**). Following quality control (**Extended Data Fig. 2b**) and doublet filtering, we recovered 55,327 high quality cells. Spatial similarity mapping using VoxHunt^39^, a tool based on *Mus musculus In Situ Hibridization* data from the Allen Brain Atlas, confirmed a hindbrain identity with a more pronounced pontine signature (**Fig. 1d**). We next generated an integrated UMAP representation of the different timepoints and performed cell-based annotation using reference datasets, including the recently published Human Neural Organoid Cell Atlas (HNOCA)^40^ and the Human Developing Brain Cell Atlas (HDBCA)^41,42^ (**Fig. 1e and Extended Data Fig. 2 c-g**). Temporal analysis revealed an initial phase of high proliferation that diminished over time (**Extended Data Fig. 3a**), as cells transitioned from pluripotent stem cells to neuroepithelium, radial glia and into distinct neuronal and glial populations that emerged by days 14 and 60, respectively (**Fig. 1f**), reflecting the natural occurring segregation of neuro- and gliogenesis phases^41^. Projection of the organoid dataset onto the HNOCA that has been annotated for neuronal lineages (**Fig. 1g**) revealed that most annotated neuronal precursor cells (NPCs), neuroblasts and neurons originated from a heterogeneous cluster spanning the hypothalamus, midbrain, and hindbrain (**Fig. 1h**), reflecting non-telencephalic neurogenesis. These neuroblasts and early neurons expressed *STMN2* and *RBFOX3* (NeuN) but lacked the telencephalic marker *FOXG1*^41^ (**Extended Data Fig. 3b**). Importantly, we observed an increased representation of neurons from the midbrain, medulla, and pontine regions, which collectively form the brainstem^43^ (**Fig. 1h**). Furthermore, neurotransmitter transporter analysis revealed a predominance of excitatory (glutamatergic) and inhibitory (GABAergic) neurons, the latter known to form synapses with DMG and promote its growth^44,45^. Smaller proportions of cholinergic and dopaminergic neurons were also detected, consistent with their distribution in the HDBCA (**Extended Data Fig. 3c,d**). In addition, immunofluorescence analysis identified Tryptophan hydroxylase 2-expressing (TPH2+) cells (**Extended Data Fig. 3e**), a key enzyme involved in serotonergic synthesis, suggesting that - although undetectable at the scRNA seq level similar to the HDBCA^41^ - this population of neurons is present. Thus, consistent with findings from the HNOCA, but also the adult brain, where hypothalamic, brainstem, and hindbrain neurons display pronounced subset heterogeneity and intermixing (referred to as Splatter neurons) compared to cortical neurons^40,46^, our organoid model mirrors this regional heterogeneity in non-telencephalic neuronal populations, with a significant enrichment in brainstem identity compared to profiles described in most HNOCA protocols.

### Pontine-medulla glial-enrichment

DMG is rooted in the glial lineage^13^, prompting us to investigate the glial composition within our organoid model. First, we showed the presence of committed astrocytes (GFAP+, AQP4+) and oligodendrocytes (OLIG2+) at the protein level (**Extended Data Fig. 3f**). At the single-cell transcriptomic level, we identified glial populations spanning pre-OPCs, OPCs, committed oligodendrocyte precursors (COPs), glioblasts, and astrocytes, offering a detailed representation of glial diversity and maturation states (**Fig. 1e**). By comparing age-matched cells of HNOCA-covered protocols, BrOs shows significant enrichment in the glial lineage, particularly glioblasts and OPCs (**Fig 1h)**. Additionally, we assessed glycolysis, an indicator of cell stress in brain organoids^40^ . Consistent with models described in the HNOCA, we observed similar glycolysis levels (**Extended Data Fig. 3g**). However, in the glial lineage, glycolysis levels were lower (**Extended Data Fig. 3h**), suggesting reduced stress and a healthier metabolic state of glial cells in our model. Moreover, OPC (referred to as oligo in the HDBCA^41^) and glioblast glial populations demonstrated a reduced number of DEGs compared to HNOCA datasets (**Extended Data Fig. 3i, Supplementary Table S3a-f**), reflecting higher transcriptional fidelity and closer alignment with primary counterparts in the HBDCA^41^. To date, no comprehensive region-wide analysis has been conducted on glial cells derived from organoids. However, the HBDCA revealed strong region-specific patterns in the glial lineage in early brain development, which could be particularly relevant for H3.3K27M-altered DMG that predominantly arises in the brainstem pontine region. Projection of the BrO datasets onto the HBDCA latent space (**Extended Data Fig. 3j, k**) and comparison with organoid protocols embedded in the HNCOA, revealed significant coverage of glial clusters present within the HBDCA. Notably, 13 out of the 19 gained clusters, as compared to models described in the HNOCA, exhibited pontine and medulla-specific identities (**Fig 1i, j, Extended Data Fig. 3j, k, Supplementary Data Table S3g**). Thus, our newly generated BrO model offers a valuable experimental framework for studying gliogenesis within the context of pontine-medulla regionality, which could prove highly relevant for modelling DMG.

### *De novo* generation of H3.3K27M-altered DMG

We next investigated whether this BrO model could be exploited to model DMG tumours. The most common H3.3K27M-defining DMG mutation ^47^, alongside typical accompanying and pons-specific tumour suppressor TP53 and platelet-derived growth factor A (PDGFRA) alterations^8,48–50^, expressing plasmids^16^ were introduced using *in situ* electroporation of developing BrOs (**Fig. 1a**). This mutation cocktail has been shown to be time-sensitive in *in utero* electroporation mouse models^14,16,17^, hence we tested different time points of electroporation between day 11 and 28. We identified day 11 as the time point most efficiently inducing tumorigenic growth (**Extended Data Fig. 4a,b, left panel, Supplementary Table S1b**), reinforcing the concept of a restricted early developmental time window for DMG transformation^14,16^. At this stage of development, we observed a dominance of radial glia (RG) and neuroepithelial stem-like cells in BrOs (**Fig. 1f**), aligning with earlier work identifying neural stem cells (NSCs) as a permissive cell state for H3.3K27M-driven neoplastic transformation^14,16,18–20^. Tracking tumour growth over two months showed that the resulting tumours display infiltrative growth (**Extended Data Fig. 4c)**. In contrast, the use of empty control plasmids resulted in only a few localized electroporated cells (**Extended Data Fig. 4b, right panel**). Whole-organoid 3D imaging on week 16 (4 months post-electroporation) with tumour colour-coded for invasion depth further confirmed a diffuse growth pattern characteristic of DMG (**Fig. 2a**). In addition, DMGOs orthotopically transplanted in immunodeficient mice were able to progress *in vivo*, demonstrating invasive growth (**Extended Data Fig. 4d**). Quantification of H3.3K27M expression, combined with dominant negative TP53 (DNTP53), and PDGFRA-D842V at the protein level showed incorporation of all three mutations into the majority of GFP-positive cells with 88%, 80%, and 76% expressing cells respectively (**Fig. 2b-d, Supplementary Data Table S1c**). These findings illustrate DMG invasive outgrowth in our guided brain organoids dependent on combined common driver mutations typically observed in patients.

**Figure 2.**
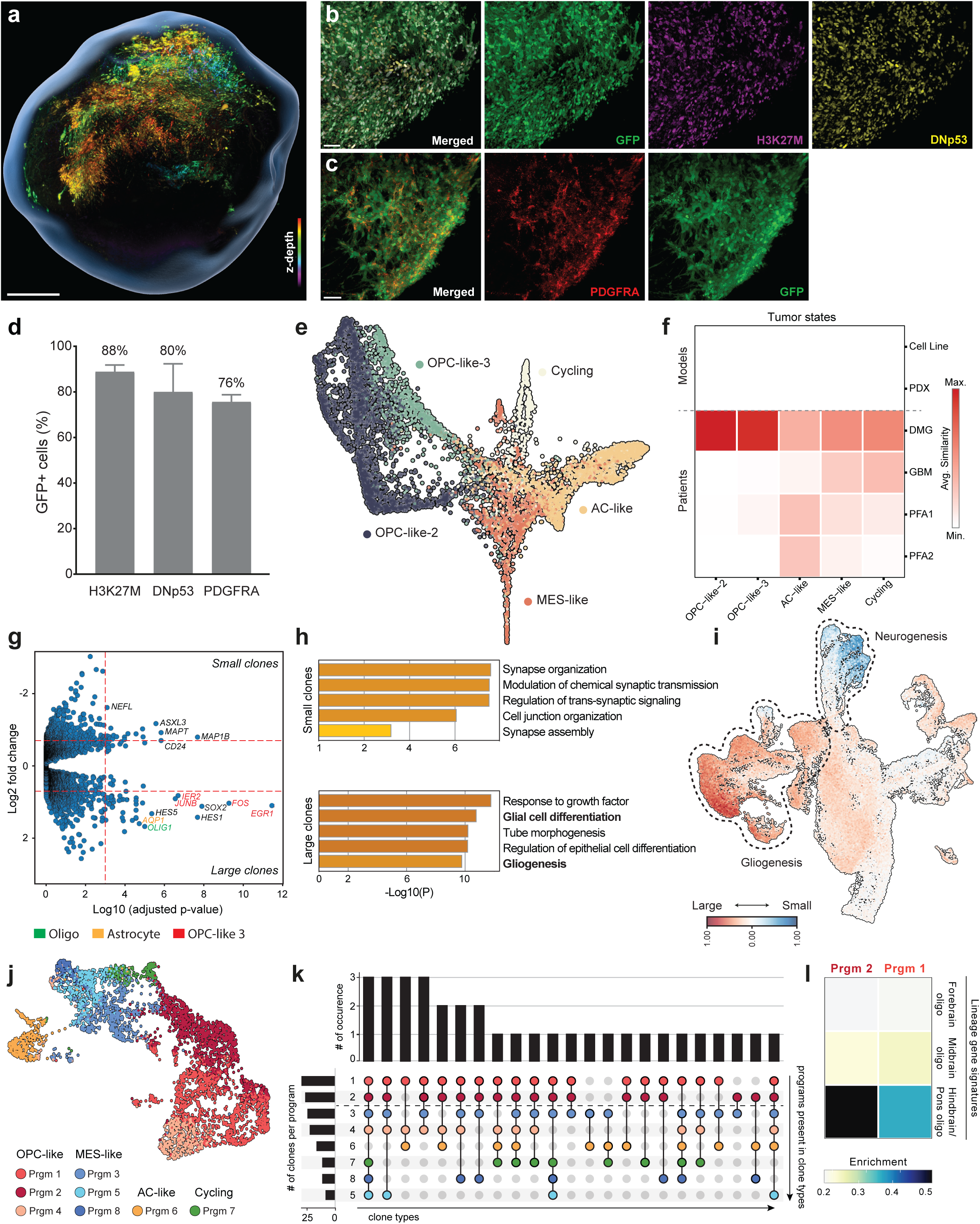
DMG tumour induction and characterization in brainstem organoids. **a**, Immunofluorescent 3D image of an intact DMGO at day 112. GFP signal colour-coded for z-depth on a rainbow scale, grey outline created by masking of Propidium Iodide fluorescence. Scale bar = 500 µm. **b, c**, Representative multispectral 3D images of tumour GFP (green), H3K27M (magenta) and DNp53 (yellow) (b) or tumour GFP (green) and PDGFRA (red) (c) in consecutive slices of a week 8 DMGO. Scale bars = 50 µm. **d**, Percentage of GFP^+^ tumour cells expressing H3K27M, DNp53, or PDGFRA detected by multispectral 3D imaging as in b and c. **e**, Integrated Force Atlas (FA) representation of DMGO tumours coloured by tumour cell state. **f**, Heatmap representation of average transcriptomic similarity between DMGO tumour cells and *in vitro* models (Cell lines and Patient-derived Xenografts, PDX), H3K27-altered DMG, Glioblastoma (GBM) and H3K27-altered Posterior Fossa Group A Ependymoma (PFA1/2, H3K27M/EZHIP-mutants) patient samples. Average similarity (colour intensity) represents an averaged prediction score of all DMGO subsetted tumour cells mapped into a merged dataset consisting of transcriptomic *in vitro* and patient datasets^11–13^. **g**, Volcano plot showing top differentially expressed genes in larger and smaller clones. **h**. METASCAPE results showing selected GO terms from the highest scoring summary GO terms for small and large clones. **i**, Presence of large (red) and small (blue) clones in the integrated UMAP representation of developing BrOs, showing a preference for gliogenesis and neurogenesis, respectively. **j**, UMAP of traced DMGO cells, coloured by their respective highest scoring cNMF program. **k**, UpSet plot displaying clonal intersection events. Only clonal families found in more than 1 cNMF module are depicted and filtered with at least 3 cells present per unique barcode. Bar plots depict the frequency of each lineage combination (top) and the number of clones that contains each program (left). Colouring of dots matches cNMF program annotation as in j. **l**, Heatmap presenting the mean cellular enrichment scores of cNMF programs 1 and 2 for forebrain-, midbrain-and hindbrain/pons oligo lineage signatures in HDBCA^41^ .

### Patient-representative DMG

To further assess the patient-representability of our *in vitro* grown tumour model, we conducted histological analysis and compared it to patient samples with the same mutational profile. This shows that H3.3K27M cells (H3K27M+) display loss in H3K27 trimethylation (H3K27me3) in both patient samples and DMGOs (**Extended Data Fig. 5a, b**), a hallmark of H3K27-altered DMG^7,47^ and again confirms invasive diffuse growth (Neurofilament (NF)+/GFAP+ tumour cells) (**Extended Data Fig. 5b**). Furthermore, our *in vitro*-grown tumours exhibited a global methylation profile closely resembling DMG, distinguishing our tumours from Glioblastoma and Posterior Fossa Ependymomas (PFA1 and 2), where the ependymomas are presenting with a similar loss of H3K27M trimethylation caused by a H3K27M mutation or EZHIP overexpression respectively^51^ (**Extended Data Fig. 5c**). We next conducted scRNA-seq profiling of sorted GFP^+^ tumour cells and after quality control filtering analysed approximately 7,000 cells from 14 DMG organoids (DMGOs) (**Fig. 2e, Extended Data Fig. 6a-b**, **Supplementary Table S2c**). The malignant state of these cells was further supported by analysis of inferred copy number variation (iCNV) from scRNA-seq data, which showed large-scale amplifications and deletions in these cells compared to healthy cells, including loss of chromosome 10 and 13 and a gain of chromosome 19q (**Extended Data Fig. 6c**). Using published DMG references^11,13^, we first annotated cancer cell states previously described for DMG, including OPC- and AC-like, as well as a MES-like cell state and a population of cycling cells. In line with the early developmental window of our model, we identified only few cells with a more mature OC-like phenotype. Importantly, we identified a major proportion of OPC-like tumour cells that resembled recently defined OPC-like-2/3 states, both described as paediatric and pons-specific pre-OPC states^13^ (**Fig. 2e, Extended Data Fig. 6d-f**). For each identified tumour state, we quantified the resemblance of DMGOs to current widely applied *in vitro* and *in vivo* models, as well as transcriptomic data from patients. Importantly, we found the highest similarity score between DMGOs and primary DMG patient material^11^, as opposed to cell lines, patient-derived xenografts (PDXs), Glioblastoma^11^ and both PFA subtypes patient material^52^ (**Fig. 2f**). Together, this highlights the ability of DMGOs to closely mimic primary DMG tumours.

### Pontine glial-specific DMG tumorigenesis

Next, we investigated the mechanisms driving tumorigenesis to identify the attributes of BrOs that appear to be critical for supporting the growth of DMG tumours. We used TrackerSeq, a PiggyBac-based genetic lineage tracing approach (**Extended Data Fig. 7a**), and analysed cancer clone dynamics at two months post-electroporation. We retrieved 167 unique barcodes from six DMGOs and two healthy BrOs (**Extended Data Fig. 7b-h, Supplementary Table S4a**) and detected individual clones spanning up to approximately 800 cells per barcode, indicative of cancerous transformation (**Extended Data Fig. 7i, j**). By comparing large versus small, traced clones (**Fig. 2g, h**) via DEG and METASCAPE analysis, we identified glial specification as a critical feature driving cancer clone expansion in contrast to the more neuronal specification enriched in smaller clones (e.g. synapse organization and modulation of chemical synaptic transmission) (**Fig. 2g, h, Supplementary Table S4b-e**). Larger clones were characterized by higher gene expression of *OLIG1*, a canonical OPC marker, but also *IER2*, *JUNB*, *FOS*, *EGR1*, previously described as key markers of the OPC-like 3 pre OPC state^13^. Interestingly, we also identified a higher expression of *AQP1*, an aquaporin previously shown to be exclusive to astrocytes in the human brainstem^53^. Furthermore, analysis of patient data^12^ revealed *AQP1* expression only in tumours located in the pons, but not in those arising from the cortex or thalamic regions (**Extended Data Fig. 7k**). These data hint towards a glial-specific tumorigenic process that is, furthermore, pontine location-dependent. This is further illustrated by genes upregulated in large DMG tumour clones mapping back to the glial lineage of our developing brainstem organoids (**Fig. 2i**), indicating that tumorigenesis is dependent on gliogenesis modelled with brainstem organoids. To confirm this experimentally, we performed *in situ* electroporation of our mutation cocktail in unguided cerebral organoids, revealing a significant reduction in tumour induction (**Extended Data Fig. 4e**). In addition, the outgrowth was non-diffuse, with almost no GFP-positive tumorigenic cells carrying the H3.3K27M mutation (**Extended Data Fig. 4f, g**). These findings illustrate that the tumorigenic potential of H3.3K27M-altered DMG is strongly dependent on the correct anatomical cellular identity that we could model with our guided brain organoids. Consensus non-negative matrix factorization (cNMF) (**Fig. 2j, Extended Data Fig. 7l, m and Supplementary Table S5a**) and lineage relationship analysis identified malignant meta-gene programs 1 and 2 to be present in the highest number of clones (30 and 26 out of 34 clones, respectively; **Fig. 2k**) and belonging to the OPC-like lineage, emphasizing the central role of this lineage in H3K27M DMG tumorigenesis^13^. More specifically, we show overlap with the pre-OPC states; OPC-like 2 (**Extended Data Fig. 7m**). In the context of human early gestation, regionally distinct gene signatures for the glial lineage have been suggested to underlie the strong region-specific occurrence pattern of glial-related diseases, such as DMG^41^. In line with this, both programs 1 and 2 specifically enrich for the hindbrain-pons oligodendrocyte precursor lineage (referred to as oligo in the HDBCA^41^), as opposed to midbrain and forebrain (**Fig. 2l, Supplementary Table S5b**). Collectively, these data highlight the critical role of pons glial specification, enriched within BrOs, in driving DMG tumorigenesis. This underscores the importance of integrating precise spatial and developmental contexts for DMG and establish this human-relevant, experimentally accessible model as an adequate tool for therapeutic evaluation.

### Modelling CAR T cell heterogeneity and functional exhaustion

Given patient-relevant tumour progression observed in DMGOs, we assessed whether they could serve as a human *in vitro* platform for preclinical evaluation of GD2 CAR T cells (**Fig. 1a**), especially because of the highly encouraging yet variable treatment outcomes of these cells in a recent first clinical trial in patients with H3K27M-mutant DMG^3,4^. By exposing untransformed BrOs to GD2 CAR T cells, we first visually inspected with brightfield imaging that the presence of GD2 CAR T cells did not affect the general health of the model (**Extended Data Fig. 8a**). Next, we confirmed GD2 target expression in DMGO (**Extended Data Fig. 8b**), as well as tumour cell killing via confocal imaging of cleaved caspase 3 in GFP^+^ tumour cells (**Fig. 3a**). Having established these experimental preconditions, we treated DMGOs four months after tumour induction by administrating CD8^+^ GD2 CAR T cells at day 0 and 7 and monitored T cell activation, measured by IFNγ secretion (**Extended Data Fig. 8c**), and tumour control (**Extended Data Fig. 8d, e**) over time. Similar to heterogenous outcomes reported in patients^3,4^, we observed an overall partial reduction in tumour burden (**Fig. 3b**) and heterogenous response rates over time and between individual DMGOs. (**Extended Data Fig. 8d, e, Supplementary Table S6a**). Since GD2 CAR T cell activation was evident by a robust IFNγ response for all treated DMGOs (**Extended Data Fig. 8c**), limited response profiles (e.g. DMGO179) are unlikely to result from a lack of antigen recognition. Importantly, therapy effects could be detected even after >1 month of treatment (**Fig. 3c**), offering advantages for modelling CAR T cell functionality *in vitro* in a manner that is representative of T cell states at the tumour site *in vivo*, including potential exhaustion profiles associated with prolonged tumour exposure. To test our model for this purpose, we sequenced over 30,000 GD2 CAR T cells retrieved from DMGOs, as well as unexposed GD2 CAR T cells. This revealed a substantial level of heterogeneity induced upon DMGO exposure (**Fig. 3d**). In GD2 CAR T cells retrieved from DMGOs, we identified 9 transcriptional states (**Fig. 3e**) that, based on combined interrogation of curated gene signatures (**Extended Data Fig. 9a**), DEGs (**Supplementary Table S6b**), DEG-associated GO terms (**Extended Data Fig. 9b-f**), expression of canonical immune effector (**Fig. 3f**) and exhaustion markers (**Fig. 3g**) and comparison to a pan-cancer infiltrating T cell (TIL) dataset that includes brain malignancies^5^ (**Extended Data Fig. 9g**), reflected different T cell activation, differentiation and effector states. For instance, we identified a GD2 CAR T cell population that, although activated (based on *HLA* gene expression) (**Extended Data Fig. 9a**), is not fully differentiating towards effector function (undifferentiated; T_UND_) (**Extended Data Fig. 9b, h**), as well as an *IL-2* responsive population (T_IL-2_) (**Extended Data Fig. 9c, i**), probably differentiating into effector T cells (**Extended Data Fig. 9i**). In addition, we observed an interferon-stimulated gene (ISG) expressing population (**Extended Data Fig. 9a**) (T_ISG_) strongly corresponding to *ISG* expressing TILs^5^ (**Extended Data Fig. 9j**) and considered an interferon-induced activation state^54,55^. Other clusters included a CAR T cell population with migrating properties and interconnectivity (T_MI_) that appears to be influenced by its neuronal environment (**Extended Data Fig. 9d)**, as well as proliferating (T_PR_) (**Extended Data Fig. 9e)** and metabolically stressed T cells (T_MS_) (**Extended Data Fig. 9a, f**). Importantly, we distinguished potential DMG-targeting effector T cell populations based on their cytotoxic profile (**Fig. 3f**) and putative level of exhaustion (**Fig. 3g**). While one of these clusters predominantly expressed *GZMK* (T_GZK_), cytotoxic T cells (T_CYT_) expressed *GZMB*, *PRF1* and *IFNG* (**Fig. 3f**). In contrast, exhausted T cells (T_EX_) displayed reduced *IFNG* and concomitant expression of immune checkpoint genes; *LAG3*, *HAVCR2*, *TIGIT*^56^ and *SELPLG*^57^, as well as the transcriptional repressor *PRDM1* associated with exhaustion^58^ (**Fig. 3g**). This demonstrates the potential advantage of prolonged treatment in DMGOs for uncovering T cell functional exhaustion, considered an actionable axis to enhance treatment outcomes^59^ and, therefore, critical to recognize during pre-clinical evaluation.

**Figure 3.**
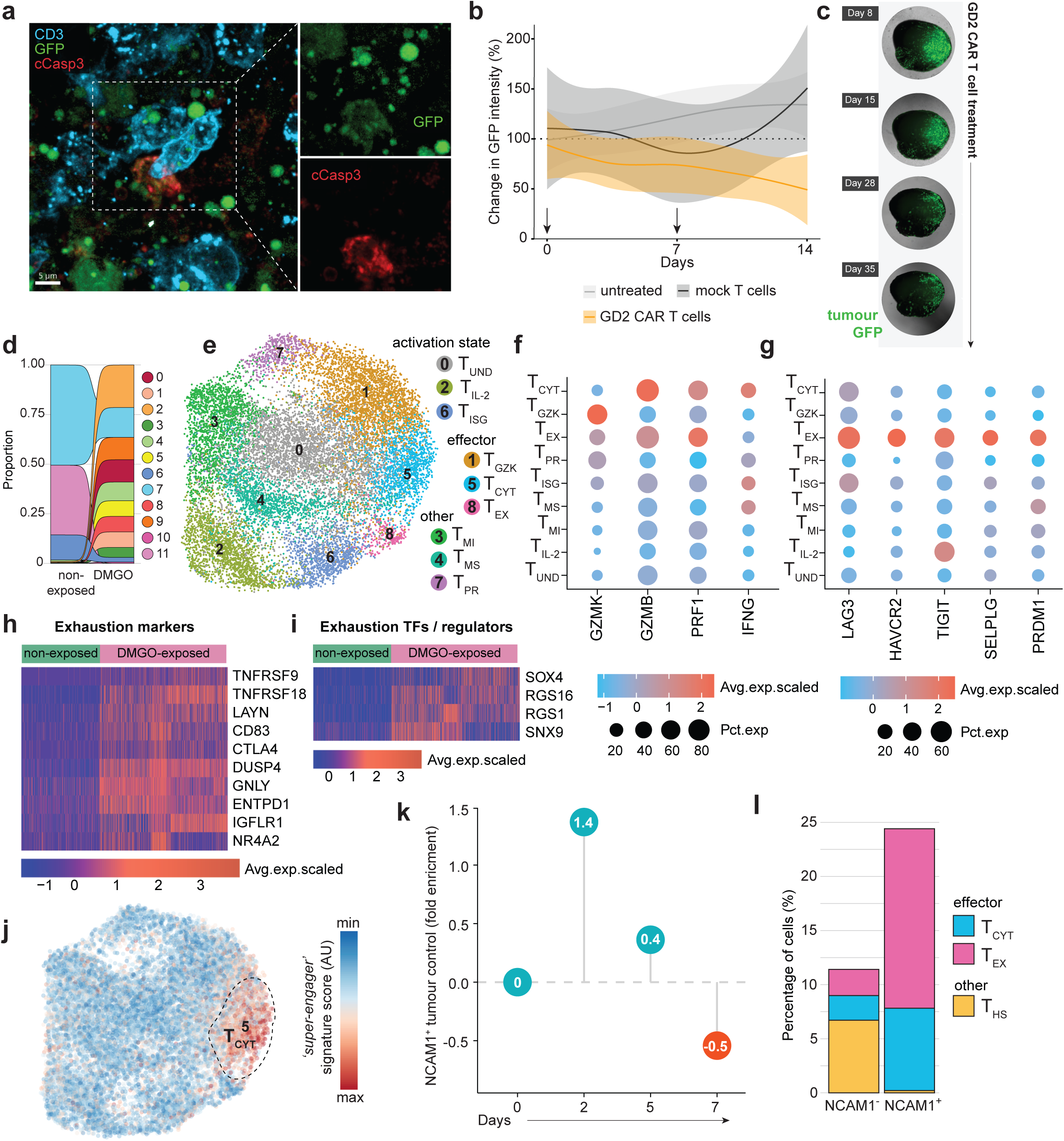
DMGOs model CAR T cell functional heterogeneity. **a**, Multispectral 3D imaging of GD2 CAR T cells (CD3; cyan), DMG tumour cells (GFP; green) and cleaved caspase-3 (cCasp3; red). Scale bar = 5 µm. **b**, GD2 CAR T cell treatment outcome measured as a relative change in tumour GFP intensity quantified by imaging compared to the start of treatment (100%). DMGOs were either left untreated (grey line, n=1), treated with mock transduced T cells (black line, n=2), or GD2 CAR T cells (orange line, n=4) and for each treatment condition a smoothed line trend between the averaged values at different timepoints was plotted using the LOESS algorithm. Shaded area reflects the 95% confidence interval, arrows indicate the timepoints of T cell administration. **c**, Representative images of the tumour GFP signal at the indicated timepoints for a DMGO subjected to prolonged GD2 CAR T cell treatment administrated at day 0, day 8 and day 15. **d**, Sankey plot illustrating the shift in the relative proportions of unbiasedly identified GD2 CAR T cell clusters before and after DMGO exposure. **e**, UMAP visualization of annotated GD2 CAR T cell clusters. **f**, Cytotoxic effector molecule and cytokine gene expression across the GD2 CAR T cell clusters. **g**, Gene expression of selected exhaustion associated receptors, ligands, and transcription factors across the GD2 CAR T cell clusters. **f**, **g**, Dot plot representing the percentage of cells expressing selected genes. Colour intensity represents the average scaled gene expression. **h**,**i**, Heatmap depicting the relative expression of exhaustion markers (h) and exhaustion associated transcription factors and functional regulators (i) in non-exposed (left) and DMGO-exposed (right) GD2 CAR T cells within the T_EX_ cluster. **j**, ‘*Super-engager*’ signature score (in arbitrary units (AU)) on a blue-to-red colour scale showing enrichment of previously identified T cell serial killer gene set from Dekkers *et al.*^67^ atop UMAP cell embeddings of the GD2 CAR T cell dataset. Dashed outline annotates the embedding of the T_CYT_ GD2 CAR T cell cluster. **k**, Dotchart depicting the fold enrichment in tumour killing by NCAM1^+^ GD2 CAR T cells over NCAM1^-^ GD2 CAR T cells quantified as the change in tumour area detected by GFP compared to the start of treatment. n=2 DMGOs per treatment condition. **l**, Percentage of cells per T_CYT_, T_EX_ and T_HS_ cluster for NCAM1^-^ GD2 CAR T cells (left) and NCAM1^+^ GD2 CAR T cells (right).

To confirm that exhaustion detected in our DMGO model reflects patient-representative T cell exhaustion at the tumour site, we compared the T_EX_ phenotype present upon DMGO exposure, to pre-exposure GD2 CAR T cells that – although alleviated by the 4-1BB endodomain - can still display exhaustion features resulting from tonic signalling^60^. Indeed, a fraction of pre-exposure GD2 CAR T cells overlapped with our T_EX_ cluster detected upon DMGO exposure (**Extended Data Figs. 10a, b**). However, separating the cells in this cluster based on experimental condition (**Extended Data Fig. 10b**) revealed that DMGO-exposed T_EX_ upregulated a wide array of additional exhaustion markers (**Fig. 3h**), as well as known TFs and functional modulators of exhaustion (**Fig. 3i**) that, importantly, include those described in patients across TIL datasets (**Supplementary Table S6c**). In addition, overlap with exhaustion markers found in the antigen-driven lymphocytic choriomeningitis virus (LCMV) mouse model of chronic infection^61,62^, as well as an *in vitro* model of CAR T cell dysfunction based on continuous antigen exposure^63^, demonstrates that the observed exhaustion profile is antigen-driven (**Supplementary Table S6c**). For *in vitro* model systems this has not yet been achieved in the context of naturally expressed tumour-antigen, only through persistent anti-CD3 and anti-CD28 antibody stimulation^64^, or by using repeated rounds of stimulation with antigen- pulsed^65^, or overexpressing^66^ tumour cell lines^63^. Thus, DMGOs model CAR T cells functional heterogeneity, including patient-representative T cell functional exhaustion.

### NCAM1 enriches for short-lived cytotoxic effector CAR T cells

In line with their strong cytotoxic profile and no signs of exhaustion, T_CYT_ showed a high degree of transcriptomic overlap with the ‘killer’ gene signature of ‘*super-engager’* engineered T cells that we recently identified to have profound tumour targeting capacity and serial killing behaviour in a short-term co-culture assay^67^ (**Fig. 3j**). Since we previously identified NCAM1 as a selection marker to enrich for this population^67^, we exploited this strategy and the DMGO prolonged treatment model (**Fig. 1a**) to further investigate the relevance of this CAR T cell functional profile in a patient-representative treatment setting. We sorted GD2 CAR T cells based on NCAM1 expression prior to DMGO treatment (**Extended Data Fig. 10c**) and compared tumour control between NCAM1^+^ and NCAM1^-^ cells (**Fig. 3k, Extended Data Fig. 10d, Supplementary Table S6d**). This demonstrated initial potent anti-tumour activity of NCAM1^+^ GD2 CAR T cells, with a 1.4-fold enrichment in tumour control over NCAM1^-^ GD2 CAR T cells at day 2. However, this enhanced potency stabilized between day 5 and 7, with NCAM1^-^ T cells displaying more gradual anti-tumour activity over time, slightly outperforming NCAM1^+^ cells by day 7 (**Fig. 3k**), in line with a higher recovery of NCAM1^-^ cells at day 14 (**Extended Data Fig. 10e**). To gain insight into potential transcriptomic profiles explaining these differential outcomes, we performed scRNA-seq of NCAM1^-^ and NCAM1^+^ GD2 CAR T cells and mapped them back to our previously identified GD2 CAR T cells signatures (**Extended Data Fig. 10f**). This revealed an additional stressed GD2 CAR T cell cluster specific to NCAM1^-^ cells (T_HS_) (**Fig. 3l** and **Extended Data Fig. 10g**) that differed from the T_MS_ cluster through high expression of heat-shock proteins (HSPs) (**Supplementary Table S6e**) and, importantly, overlaps with the stress response state identified in patient TILs that associates with immunotherapy resistance^5^ (**Extended Data Fig. 10h**). Further aligning with the initially enhanced tumour control observed in **Fig. 3k**, NCAM1^+^ cells show a 3.3-fold enrichment in T_CYT_ compared to sorted NCAM1^-^ cells (**Fig. 3l**). However, in line with poor persistence of the cells (**Extended Data Fig. 10e**), NCAM1^+^ T cells are additionally enriched for T_EX_ (**Fig. 3l**), explaining their reduced performance over time (**Fig. 3k**). Together, this identifies NCAM1^+^ cells as the most potent tumour-targeting, yet short-lived effector GD2 CAR T cell population and offers proof-of-concept for cell selection as a means to narrow CAR T cel functional heterogeneity prior to patient administration.

### Microglia-enriched DMGO microenvironment

The upregulation of features associated with tissue-residency^68^, including the canonical marker CD103 (ITGAE) used to identify tissue-resident T cells^69,70^ (**Extended Data Fig. 10i, Supplementary Table S6c**) underscores the capacity of DMGOs to model T cell performance within tissue. While this could lead to new strategies to enhance tumour trafficking and tissue residency sought after for improving CAR T cell performance^71^, the immune myeloid compartment, also critically influencing T cell responses, is for now not included in this environment. Therefore, we sought to further enrich the complexity of our DMGO model by including microglia, a key compartment of the DMG tumour microenvironment^13^. To achieve this, we generated primitive macrophage progenitors (PMPs) from human stem cells, which have been previously shown to differentiate into mature microglia in mouse brains^72^, human midbrain organoids^73^ and in co-culture with neurons^74^. Similarly, PMPs integrated into our brainstem organoid model and showed a ramified morphology associated with homeostatic microglia^75^ within 7 days of integration (**Extended Data Fig. 11a, b**). Confirming functional maturation, the cells displayed typical microglia functionality, migrating to sites of myelin injection (**Extended Data Fig. 11c, Supplementary Movie S1**) and removing myelin through phagocytosis^76,77^ (**Extended Data Fig. 11d, Supplementary Movie S2**). Furthermore, three weeks after incorporation in brainstem organoids, above 80% of cells expressed the microglia-specific marker P2RY12 at the protein level and scRNA-seq analysis demonstrated increased expression of microglia-specific TFs^78,79^, indicating their maturation into microglia (**Fig. 4a-c**). Additionally, the cells resembled an adult state when referenced against microglia developmental programs identified in mice^80^, further validating microglia maturation (**Extended Data Fig. 11e**). Comparison with a myeloid cell reference dataset from DMG patient tumours^6^ further confirmed microglia as opposed to macrophage identity (**Fig. 4d**).

**Figure 4.**
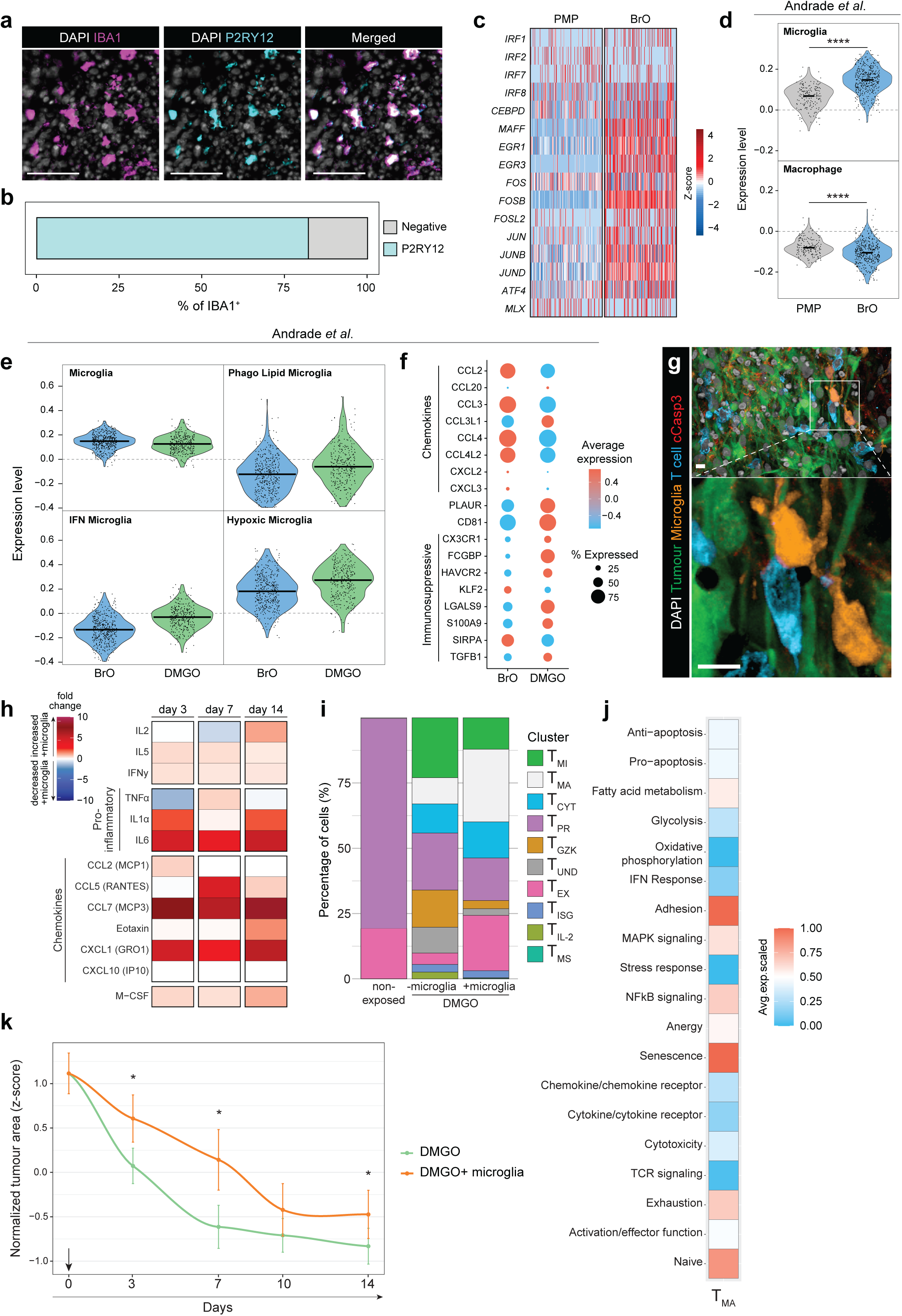
Microglia integration and impact on the GD2 CAR T cell response. **a**, Immunofluorescent 2D images of BrO with microglia integrated for 3 weeks, labelled for DAPI (white), IBA1 (magenta) and P2RY12 (cyan). Scale bar = 50 µm. **b**, Quantification of percentage of P2RY12^+^ cells of total IBA1^+^ microglia in a BrO slice. **c**, Heatmap depicting the relative expression of microglia-associated transcription factors in PMP (left) and BrO-derived microglia 3 weeks after integration (right). **d**, Violin plot showing the expression level of microglia and macrophage gene signatures from Andrade *et al*.^6^ in PMP (left) and microglia derived from BrO (right). **e,** Violin plots showing expression levels of DMG-associated microglia states from Andrade *et al*.^6^ in microglia derived from BrO (blue) or DMGO (green). **d**, **e**, ****p < 0.0001, two-tailed t-test. **f,** Dotplot showing the relative expression of selected chemokines and genes associated with immunosuppression from Andrade *et al*.^6^ in microglia derived from BrO (left) or DMGO (right). **g**, Immunofluorescent 3D images of a 200 µm thick DMGO slice containing microglia, 1 week after start of GD2 CAR T cell treatment. Cells are labelled for DAPI (white), GFP^+^ DMG tumour cells (green), IBA1^+^ microglia (orange), CD3^+^ T cells (cyan) and cleaved caspase-3 (cCasp3; red). White insert indicates zoom area displayed below. Scale bar = 10 µm. **h**, Heatmap depicting the fold change in concentration of selected cytokines, chemokines and growth factors of DMGOs containing microglia normalized against no microglia at day 3, day 7 and day 14 after GD2 CAR T cell addition. n > 3 DMGOs. **i**, Percentage of cells within GD2 CAR T cell clusters, including a new microglia-affected cluster (T_MA_), for non-exposed GD2 CAR T cells (left), or GD2 CAR T cells retrieved from DMGO without (middle) or with integrated microglia (right). **j**, Heatmap highlighting the average scaled expression of curated gene signatures from Chu *et al*^5^. in the T_MA_ GD2 CAR T cell cluster. **k**, Reduction in tumour area (normalized z-score per DMGO relative to timepoint 0) after addition of GD2 CAR T cells in DMGO without (green) or with integrated microglia (orange). Statistical analysis at each timepoint was performed using a linear mixed-effects model, accounting for experimental and organoid variation (t3; p=0.0165, t7; p=0.0219, t10; p=0.0841 and t14; p= 0.0434). Arrow indicates the timepoint of GD2 CAR T cell administration.

Importantly, microglia incorporation in tumour-bearing DMGOs led to the acquisition of DMG-associated diverse functional phenotypes recently described^6^, including an interferon-activated (IFN), phago lipid and hypoxic state (**Fig. 4e**). Moreover, while microglia from brainstem organoids displayed GO terms related to neuronal surveillance and protein production, microglia from DMGOs were characterized by GO terms related to antigen presentation and immune responses, such as processing and presentation of peptide antigen via MHC-II and response to type I IFN, which have been previously described to be upregulated in DMG-associated microglia/macrophages^81^ (**Extended Data Fig. 11f, Supplementary Table S6f**). In addition, these DMG-specific microglia states were accompanied by reduced chemokine expression and upregulation of genes associated with an immunosuppressive profile, in line with what has been shown in patient data^6^ (**Fig. 4f**). This was confirmed by protein expression of CD163, associated with an anti-inflammatory state, and SPP1, associated with immunosuppressive lipid-laden macrophages^82^, in microglia from DMGOs (**Extended Data Fig. 11g, h**). Together, this demonstrates that under the influence of the appropriate neuronal environment adequately modelled in brainstem organoids, PMPs differentiate into mature functional microglia that, importantly, in the presence of DMG tumour cells resemble a DMG-specific immunosuppressive state.

### Microglia impact GD2 CAR T cell therapy responses

Correlative clinical trial data suggest that a rise in the immunosuppressive myeloid compartment might coincide with unfavourable GD2 CAR T cell treatment outcomes^4^. In addition, myeloid cells, including microglia, are considered mediators of CAR T cell induced toxicity^83^. To address this experimentally, we performed GD2 CAR T cell treatment in DMGOs with integrated microglia. Confocal imaging showed interactions between GD2 CAR T cells and microglia within tumours (**Fig. 4g**). Moreover, we observed increased cytokine secretion in the presence of both GD2 CAR T cells and integrated microglia (**Fig. 4h**). This included upregulated chemokines related to myeloid cell chemotaxis, for instance, MCP3 and CXCL1, as well as myeloid cell associated growth factors, e.g. M-CSF. In addition, we observed an increase in key pro-inflammatory cytokines implicated in CAR T cell (neuro-)toxicity; IL-1α and IL-6 that are routinely blocked in the clinic to manage toxicity^3^. This indicates that microglia-integrated DMGOs might offer a suitable experimental model to delineate CAR T cell microglia interaction involved in adverse treatment effects. Furthermore, when we analysed DMGO-induced CAR T cell transcriptional heterogeneity in the presence or absence of microglia (**Fig. 4i**), we observed an increase in the relative proportion of the exhausted cluster (T_EX_) and identified a new population of microglia-affected GD2 CAR T cells (T_MA_) that upon closer inspection predominantly mapped to undifferentiated TIL populations from the Chu *et al*. pan-cancer atlas (naive and tissue-resident memory cells)^5^ (**Extended Data Fig. 11j**). To further confirm the low effector profile of these cells, we plotted the expression of curated gene signatures from the same resource^5^ for this T_MA_ population relative to the other identified GD2 CAR T cell clusters (**Fig. 4j, Extended Data Fig. 11k**). Among others, this revealed low expression of signatures related to T cell activation and effector function, including cytotoxicity, while expression of genes related to senescence was high. Thus, the presence of microglia shifts the transcriptional profile of GD2 CAR T cells towards reduced effector function. In line with this, when we monitored CAR T cell mediated tumour control it was significantly reduced in DMGOs with integrated microglia (**Fig. 4k**). Together, this shows that microglia can successfully be integrated in the DMGO model, reflecting a patient-representative phenotype that can subsequently be interrogated for its impact on CAR T cell functionality and toxicity in this experimentally accessible and patient-representative model. This critical application for the first time demonstrates a direct impact of the presence of microglia on CAR T cell functional profiles, culminating in reduced tumour control.

## Discussion

Here, we show that guided differentiation of brain organoids using a combination of FGF4 and retinoic acid generates brainstem-patterned organoids (BrOs), enabling both spatial and temporal modelling of the developing foetal brainstem, including pontine-specific features, the regional origin and essential niche for H3.3K27M-altered DMG formation. We exploited these critical features to successfully established a *de novo* DMG model that faithfully recapitulates key features of patient tumours and demonstrated the central role of hindbrain-pons OPC lineages in DMG tumour progression.

Uncovering DMG’s therapeutic vulnerabilities depends on faithfully recapitulating the patient’s pathobiology in a preclinical setting. In this context, our study conceptually expands the existing spectrum of human-centred models. While patient-derived organoids (PDOs) for DMG are starting to be used for drug assessment^67,84–87^, our model provides an additional solution to generate *in vitro* tumours for this rare and deadly disease that affects only 1,000 patients annually and for which patient material is limited. Furthermore, the use of iPSCs as a cell source makes patient-specific modelling a future possibility, creating an *in vitro* system for personalized drug evaluation, from which results could be directly compared to clinical outcomes. Importantly, our model enables prolonged treatment studies, within the context of the location-specific neuronal enriched microenvironment, offering a critical tool for advancing therapeutic research. Prolonged treatment of DMGOs reflects variable treatment outcomes as observed in patients^3,4,88^ and demonstrates a high level of CAR T cell functional heterogeneity. From this heterogeneity, we identified the most potent, yet short-lived, CAR T cell population and validated a means to enrich for these cells, offering a potential approach to optimize therapy composition^89^. In addition, this model could be further leveraged in the future to reveal how CAR T cells might modulate cancer cell states to identify mechanisms of acquired treatment resistance that could be complementary targeted to enhance clinical efficacy. However, given the recognition that the tumour microenvironment can significantly impact treatment response, we focussed on implementing a critical cellular player of the local tumour immune microenvironment, by integrating microglia derived from human embryonic stem cells^79^.

In line with patient-representative tumour cell states observed in DMGOs, microglia differentiated into DMG-specific and largely immunosuppressive profiles^6^. This allowed us to for the first time experimentally address the impact of these cells on CAR T cell functional heterogeneity, revealing enhanced functional exhaustion, but also identifying a new microglia-affected population (T_MA_) with stalled differentiation and a low effector profile. This shift toward dysfunctional states correlates with reduced tumour control in the presence of microglia, offering an experimental framework to develop therapeutic strategies that mitigate microglia-induced resistance and enhance antitumor activity. In addition to preclinical development of combinatorial therapies focused on myeloid cell suppressive features^90^, our framework allows to specifically tailor these efforts to CAR T cell biology. While the occurrence of T cell functional exhaustion could potentially be mitigated by combined immune checkpoint inhibition, therapeutic strategies counteracting the newly identified T_MA_ population require further investigation of the underlying mechanism involved. Our imaging data revealing direct interactions between microglia and CAR T cells, suggest that a more thorough understanding of the nature of these interactions and the signalling pathways involved could be a critical starting point for this future work. Similar strategies could be followed to experimentally assess the influence of microglia on tumour cell phenotypes. For instance, prior work identified a link between mesenchymal tumour cell states and tumour-associated macrophages, although more pronounced in older patients^13^.

Altogether, we generated a bona fide human organoid model for DMG with critical applications towards understanding CAR T cell functionality in the context of the tumour microenvironment that could aid further therapy development for this detrimental disease.

