## Supplementary Material for "*De novo* H3.3K27M-altered Diffuse Midline Glioma in human brainstem organoids to dissect GD2 CAR T cell function"

#### Supplementary Figure legends

**Extended Data Figure 1. Brainstem organoid specification and reproducibility. a**, Overview of patterning approach and corresponding representative brightfield images of brainstem organoids over time. Scale bar of single organoids and multiple organoids is 500 µm and 2 mm, respectively. **b**, Schematic representation of a human foetal brain in gestational week (GW) 05 with indicated morphogens influencing the differentiation of hindbrain rhombomeres (r) and their HOX gene code respectively. **c**, Heatmap showing relative bulk RNA expression of homeobox (*HOX*) genes at week 02 and 03 depicted as log2 fold change normalized to week 01. Data from 3 independent batches with 3 pooled organoids per batch. **d**, Boxplot representation of Spearman's rank coefficient between different organoid batches from week 02 to 12. **e**, PCA of organoids at different timepoints from week 01 to 12 (grey scale) and derived from hESCs (circles) or iPSCs (square).

**Extended Data Figure 2. Quality control and integration of brainstem organoids time course scRNA seq data. a**, Unintegrated UMAP representation of developing brainstem organoids, coloured by collection day. Upper left corner depicts the contribution of each day to the final dataset. **b**, Violin plots showing the number of genes, number of counts, percentage of mitochondrial genes and percentage of ribosomal genes after filtering on a sample-by-sample basis. **c**, scIB integration benchmarking of different assessed integration methods, showing metrics for preservation of biological variation and batch correction. **d**, scPoli latent embedding of brainstem organoid cells, coloured by timepoint (top) and snapseed annotation (bottom). **e**, scPoli integrated UMAP of brainstem organoids, coloured by timepoint. **f**, scPoli integrated UMAP representation showing three age bins, early (day 5 – day 20), mid (day 30 – day 60) and late (day 90 – day 120). **g**, Matrixplot of scPoli (He *et al.*[40], HNOCA) or scANVI (Braun *et al.*[41], HDBCA) mediated label transfer annotation compared to the final annotation.

**Extended Data Figure 3. scRNA seq and 3D imaging characterization of brainstem organoids.** **a**, Area plots showing relative ratios of the cell cycle phases over the different timepoints. **b**. Dotplot showing the expression of selected markers for each cell type annotation, coloured by their respective cell class. *FOXG1* is represented in bold, indicating absence of expression. **c**, Annotation of neurotransmitter transporters (NTT) for cells assigned as neuroblast or neuron. **d**. Comparison of NTT found in the brainstem organoid dataset and the HDBCA pons. **e**. 3D confocal image of a 200 µm thick organoid slice at day 110 labelled for TUBB3 (orange) and TPH2 (blue). White insert indicates zoom area displayed on the right. Overview image and zoom scale bars 250 µm and 25 µm, respectively. **f**, Representative optical section of immunofluorescent 3D imaging of a 200 µm thick organoid slice at week 16 labelled for neurofilament (NF, white), GFAP (red-to-white gradient) and AQP4 (green) (left) or for DAPI (white) and OLIG2 (red) (right). Scale bars = 50 µm. **g**, Glycolysis scores over the HNOCA and

BrO datasets. Dashed line represents the mean score over all HNOCA datasets. **h**, Glycolysis scores for the OPC (left) and Glioblast (right) lineages in the HNOCA and brainstem organoids (BrOs), statistically analysed using a permutation test. **i**, DEG analysis comparing Oligo (left) and Glioblast (right) from the HNOCA and brainstem organoids (BrOs) to the HDBCA counterparts. **j**, UMAP of the HDBCA[41], coloured by the brainstem organoid presence scores. **k**, Proportional brainstem organoid presence scores for the HDBCA[41], showing the ratio of cells being represented in brainstem organoids (BrOs) per annotated cell class.

**Extended Data Figure 4. Tumour induction efficiency, outgrow, and mutational phenotype compared to unguided neural organoids.** **a**, Stacked bar plot quantifying electroporation efficiency (light grey columns) and tumour induction (dark grey columns) in brainstem organoids tested at various timepoints ranging from day 11 to day 28. ns = not significant, **p < 0.01, two tailed independent t-test. Mean (+S.E.M) from n=23-35 individual organoids per timepoint from 9 independent batches. **b**, Representative image of GFP expression (left; tumour-inducing mix) or control (right; PiggyBac backbone including CAG-mVenus) as a measure of tumour outgrowth at week 6. Scale bars = 1 mm. n = 139 organoids from 9 independent batches. **c**, Representative images of tumorigenic outgrowth of the same DMGO at week 4, 6 and 8. Scale bars = 500 µm. White arrowheads depict invasive and diffuse patterns. **d**, 3D confocal image of a 300 µm brain section of a DMGO-transplanted NSG mouse labelled for CD31 (red), tumour-GFP (green) and DAPI (grey). Scale bar = 500 µm. **e**, Stacked bar plot quantifying electroporation efficacy (light grey columns) and tumour induction (dark grey columns) for guided brainstem organoids as compared to unguided neural organoids at day 11. ns = not significant, *p < 0.05, two-tailed independent t-test. Mean (+ S.E.M) from n=3 independent experiments with a total of >35 individual organoids for each condition. **f**, Representative images of tumorigenic outgrowth (GFP; green) at week 4 and 8 for unguided neural organoids. Scale bars = 500 µm. n = 36 unguided organoids from 3 independent batches. **g**, Representative multispectral 3D images of tumour GFP (green), H3K27M (magenta) and DNp53 (yellow) in an unguided neural organoid at week 8. Stacked bar plots represent summarized data, see Methods for details. Individual values of each replicate are provided in Supplementary Table S1b.

**Extended Data Figure 5. Pathohistological and methylation profile comparison of DMGOs to primary DMG.**

**a**, Routine histopathological characterization of a representative DMG tumour sample harbouring *H3.3K27M*, *TP53* and *PDGFRA* mutations. Top panels; staining for H3K27M and H3K27me3. Bottom panels; Haematoxylin and eosin (HE), glial fibrillary acidic protein (GFAP) and neurofilament (NF). **b**, Routine histopathological characterization of a representative DMGO at day 120. Top panels; staining for H3K27M and H3K27me3. Bottom panels; Haematoxylin and eosin (HE), glial fibrillary acidic protein (GFAP) and neurofilament (NF). **c**, Methylation profile of DMGO (pooled sample consisting of 3 independent replicates) compared to DMG or resembling tumour types (GBM or EPN-PFA).

**Extended Data Figure 6. Quality control, processing and analysis of DMGO scRNA seq data.**

**a**, Violin plots showing the number of genes, number of counts, percentage mitochondrial genes and percentage of ribosomal genes after filtering on a sample-by-sample basis. **b**, Unintegrated UMAP representation of DMGO cells, coloured by sequencing batch. **c**, CNV profile of DMGO cells showing chromosomal aberrations compared to healthy cells from the BrO time course data shown in Figure 1e. Orange depicts tumour and green represents healthy cells. **d**, Dotplot showing marker gene expression for different tumour states, colour-coded for their respective annotation. **e**, Matrixplot with the obtained tumour state annotations compared to Jessa *et al.*[12] and Liu *et al.*[13]. **f**, Mapping scores of DMGO cells in Liu *et al.*[13], showing high presence of the annotated cell states, except for cycling cells.

**Extended Data Figure 7. Barcode representation after quality control and filtering and cNMF program annotation.** **a**, Schematic representation of the applied approach for simultaneously recovering transcriptomic information, HTO hashtags and lineage barcodes from DMGOs on a single cell level. A nested PCR strategy was applied for the TrackerSeq barcode. **b-f**, Quality control assessment and filters used to select barcodes from experimental replicate 1 (top) and experimental replicate 2 (bottom). Histogram depicting the total number of UMI counts prior to any filtering (b). Histogram displaying read counts and the cut-off (red dashed line) set as a minimum total read count of log$10^2$ and log$10^3$ for experimental replicate 1 and 2, respectively (c). Scatter plot depicting read counts plotted against UMI counts and the applied threshold for minimum total read counts indicated (red dashed line) (d). Histogram depicting the mean oversequence per barcode and thresholding applied (dashed blue line) (e). Scatter plot depicting read counts plotted against max mean oversequence showing both thresholds applied (f). **g**, UMAP embedding of lineage-traced DMGOs and BrO controls. Cells are coloured according to unique lineage barcodes. Cells without barcodes are presented in grey. **h**, Representation of each DMGO and BrO controls in the final UMAP embedding. **i**, Bargraph depicting the total number of cells for each clonal barcode after applying the filtering of >3 cells per clonal family. **j**, Pie charts of the relative size of each recovered clonal family among all barcoded cells per sample used for the large versus small clone comparison. Percentage is depicted for clonal families that are equal to, or above 20%, which are defined as large clones. **k**, Dotplot showing normalized expression of *AQP1* and *AQP4* in DMG cells from different locations from Jessa *et al.*[12]. In contrast to *AQP1*, *AQP4* – a canonical AC-like marker – was present in DMG tumours across all locations. **l**. Module scores of gene programs as derived from cNMF projected onto the UMAP of lineage-traced cells. **m**, Heatmap of Jaccard index scores, indicating the overlap of the cNMF derived programs to tumour state annotations.

**Extended Data Figure 8. DMGO GD2 expression and CAR T cell mediated tumour control.** **a**, Representative brightfield images at day 14 of untransformed brainstem organoids (BrOs) left untreated (n=8) or exposed to GD2 CAR T cells (n=8) at day 0 and 7. **b**, DMGO tumour cell GD2 expression (orange) analysed by flow cytometry compared to an unstained control (black). **c, d**, IFNγ levels measured in the culture supernatant (c) and GD2 CAR T cell treatment outcome measured as tumour GFP intensity relative to the start of treatment (day 0, 100%) with a smoothed line trend plotted between the values at different timepoints for each DMGO using the LOESS algorithm (d). DMGOs were either left untreated (grey line, n=1), treated with mock transduced T cells (black lines, n=2), or GD2 CAR T cells (orange lines, n=4). Arrows indicate the timepoints of T cell administration. **e**, Images of tumour GFP signal on day 0, 7, 10 and 14 for an untreated DMGO (n=1) and DMGOs treated with mock transduced T cells (n=2), or GD2 GAR T cells (n=4). GD2 CAR T cells and mock transduced T cells were administrated at day 0 and 7. **c-e**, DMGOxxx indicates individual DMGO sample ID for reference across data figures.

**Extended Data Figure 9. Key marker genes, GO terms and reference data projection of GD2 CAR T cell clusters. a**, Dot plot showing key marker gene expression (selected from the top 20 DEGs) across the GD2 CAR T cell clusters. Dot size is proportional to the percentage of cells expressing a gene and colour intensity to the average scaled gene expression. Grid colours highlight genes that are closely related in function; *HLA* genes (green), metabolic stress-related genes (red) and ISGs (blue). **b-f**, Selected significant GO terms associated with the DEGs of the $T_{UND}$ (b), $T_{IL\text{-}2}$ (c), $T_{MI}$ (d), $T_{PR}$ (e) and $T_{MS}$ (f) GD2 CAR T cell clusters. **g**, UMAP visualization of the $CD8^{+}$ TIL clusters from the Chu *et al.* pan-cancer atlas[5] used as a reference dataset. Annotated clusters are highlighted because of their overlap with, or use in defining the GD2 CAR T cell clusters. **h-j**, Marker gene signatures (DEG analysis adjusted p-value $< 0.00001$) of the $T_{UND}$ (h), $T_{IL\text{-}2}$ (i) and $T_{ISG}$ (j) GD2 CAR T cell clusters projected onto the $CD8^{+}$ TIL dataset from g.

**Extended Data Figure 10. Non-exposed, DMGO-exposed and NCAM1 subset characterization of GD2 CAR T cells. a**, UMAP representation of the integrated GD2 CAR T cell dataset, illustrating the distribution of non-exposed GD2 CAR T cells. **b**, UMAP embedding of DMGO-exposed (pink) and non-exposed (green) GD2 CAR T cells within the $T_{EX}$ cluster. **c**, Applied gating strategy (top panels) and obtained purity (bottom panels) of sorted $NCAM1^{-}$ and $NCAM1^{+}$ GD2 CAR T cells. **d**, Representative images of tumour control measured by GFP imaging in DMGOs treated with sorted $NCAM1^{-}$ (top) or $NCAM1^{+}$ (bottom) GD2 CAR T cells. **e**, Number of $NCAM1^{-}$ and $NCAM1^{+}$ GD2 CAR T cells retrieved from each DMGO sample (n=2) after two weeks of treatment. **f**, Proportion of cells within GD2 CAR T cell clusters, including those identified in Fig. 3e, a cluster enriched in non-exposed cells, and the $NCAM1^{-}$ specific $T_{HS}$ cluster. Data is shown separately for $NCAM1^{-}$ GD2 CAR T cells (left) and $NCAM1^{+}$ GD2 CAR

T cells (right). **g**, Selected significant GO terms associated with the DEGs of the $T_{HS}$ NCAM1$^-$ specific GD2 CAR T cell cluster. **h**, Upregulated marker gene signature (DEG analysis adjusted p-value < 0.05; avg_log2FC>0) of the $T_{HS}$ GD2 CAR T cell cluster projected onto the pan-cancer CD8$^+$ TIL dataset from Chu *et al.*[5]. **i**, Gene expression of tissue resident markers in non-exposed (top) and DMGO-exposed (bottom) GD2 CAR T cells within the $T_{EX}$ cluster. Dot plot representing the percentage of cells expressing selected genes. Colour intensity represents the average scaled gene expression.

**Extended Data Figure 11. PMP integration and microglia characterization in brainstem organoids and DMGOs.**

**a**, Schematic overview of PMP integration in BrOs and DMGOs and treatment with GD2 CAR T cells. **b,** Brightfield images of BrOs with mScarlet$^+$ PMPs (white) 3 or 7 days after integration. White inserts indicate zoom area displayed on the right. Overview image and zoom scale bars 500 µm and 50 µm, respectively. **c-d,** Live 3D imaging of microglia (orange) and CFSE-labelled myelin debris (green), showing homing of microglia to sites of myelin debris injection (c) and phagocytosis of myelin debris (d). Scale bar = 50 µm (c) or 10 µm (d). **e**, Heatmap depicting representation of microglia developmental stages from Matcovitch-Natan *et al.*[80] in PMP or microglia derived from BrOs or DMGOs. **f**, Selected significant GO terms associated with DEGs in microglia derived from BrOs (left) or DMGOs (right). **g**, Immunofluorescent 2D images of DMGO with microglia, 3 weeks after integration, labelled for DAPI (white), IBA1 (green), SPP1 (magenta) and CD163 (blue). Scale bar = 50 µm. **h**, Quantification of the percentage of CD163$^+$ (left) or SPP1$^+$ cells (right) in IBA1$^+$ microglia in DMGOs (n = 3 DMGO slices). **i,** Gating strategy used to sort mScarlet$^+$ PMP/microglia and CD3$^+$ GD2 CAR T cells for scRNA-seq. **j**, Bargraph depicting proportion of the $T_{MA}$ GD2 CAR T cells matched to Chu *et al*. identified pan-cancer CD8 TIL subsets[5]. **k**, Heatmap depicting the average scaled expression of curated gene signatures from Chu *et al*[5] across the GD2 CAR T cell clusters.

**Supplementary Video 1. Microglia home to sites of myelin debris injection.**

Live 3D imaging of microglia (orange) and CFSE-labelled myelin debris (green). Areas of myelin debris injection are outlined by white dashed lines. Comparison of microglia homing at 0 and 14.5 hours provided at the end.

**Supplementary Video 2. Microglia phagocytose myelin debris.**

Live 3D imaging of microglia (orange) and CFSE-labelled myelin debris (green). White arrows indicate myelin debris that is phagocytosed by microglia outlined by white dashed lines.

**Supplementary Table S1.** Overview of organoids used in order of appearance in the manuscript (**a**), electroporation efficiency data, including statistical analysis (**b**) and number of tumour cells positive for H3K27M, p53, and PDGFRA, including statistical analysis (**c**).

**Supplementary Table S2.** Metadata information for bulk (**a**) and single cell sequencing performed on BrOs (**b**), DMGOs (**c**) GD2 CAR T cells (**d**), and microglia (**e**).

**Supplementary Table S3.** DEGs from single cell sequencing of Glioblasts and OPC from BrO, HNOCA, and HDBCA datasets (**a-f**) and mean gains for each HDBCA cluster in the BrO model (**g**).

**Supplementary Table S4.** Recovered barcodes per DMGO sample (**a**) and DEGs (**b,d**) and selected GO terms (**c,e**) for large (**b**,**c**) versus small clones (**d**,**e**).

**Supplementary Table S5.** Gene lists derived from consensus non-negative matrix factorization (cNMF) (**a**) and oligo lineage signatures from Braun *et al.*[41] (**b**).

**Supplementary Table S6.** Tumour GFP intensity related to Fig. 3b (**a**), DEGs from the GD2 CAR T clusters identified in Fig. 3e (**b**), selected upregulated DEGs in DMGO-exposed compared to non-exposed $T_{EX}$ (**c**), fold enrichment in tumour control for $NCAM1^{+}$ over $NCAM1^{-}$ GD2 CAR T cells related to Fig. 3k (**d**), top 10 upregulated DEGs in the $NCAM1^{-}$ specific $T_{HS}$ GD2 CAR T cell cluster (**e**), DEGs in microglia derived from DMGO compared to BrO (**f**).

**Supplementary Table S7.** List of primary and secondary antibodies and dilutions used for immunolabeling (**a**).

**Supplementary Discussion**

Here, we offer a new guided brainstem-regionalized organoid (BrO) model that we thoroughly benchmarked to the latest single-cell human organoid and brain atlases[40–42], not only focusing on the neuronal population, but also glial lineages, to offer a well-characterized model for studying neurodevelopmental processes related to brainstem biology and associated disorders[91]. However, it should be noted that our model shares general and well-known limitations of neural organoids, including a lack of vasculature, hypoxia, diffusion-limited necrotic core formation[23,92] and cellular stress[93], all of which can to some extend impair neural specification and viability. In addition, because the model is human embryonic stem cell (hESC)-derived, it is limited in recapitulating postnatal tissue and potentially reflects a developmental stage equivalent to the second trimester, as reported for other neural organoids[40,94]. While this developmental window for inducing DMG tumorigenesis aligns with previous murine[14–16,95] and *in vitro* studies[18–20] that point to an early embryonic period in which NSCs are most permissive to H3.3K27M-driven transformation, it has also been shown that H3.3K27M retains tumorigenic potential beyond postnatal gliogenesis (P0/P1)[15]. Although impossible to confirm in patients, this suggests temporal flexibility in disease initiation that we are not replicating with our DMG organoid (DMGO) model. Furthermore, this might carry consequences for accurately modelling the in-patient neuronal microenvironment, which will need to be addressed in future studies.

Electroporation of H3.3K27M and accompanying driver mutations in brainstem-regionalized organoids compared to unguided neural organoids and the application of a genetic lineage tracing approach, uncovered an essential role for the pontine glial lineage, and in particular OPCs, in fuelling DMG progression. However, this non-targeted electroporation strategy does not allow to directly address the cell-of-origin. Precise targeting of permissive cell states (e.g. $NES^+$, $OLIG2^+$) will be required to further dissect the cellular origin, regulatory trajectories, and context-specific vulnerabilities driving H3.3K27M-associated tumorigenesis in a human-relevant setting. Finally, in the currently presented model we do not capture complex intraregional interactions, which are considered especially relevant for the pons that provides a key relay between the forebrain and motor/sensory pathways. The incorporation of assembloid methodology[96–98], in between cortical organoids and BrOs/DMGOs, for example, could enhance neuronal health and lineage diversity, as well as enable the modelling of DMG cross-regional invasion and progression depending on neural secretion[45,99–101] and activity[44,102–104].

While further advancements of DMGO complexity could, thus, address current limitations, we also show that these *de novo* DMG organoids are compatible with prolonged GD2 CAR T cell treatment, mimicking both treatment outcomes[3,4] and T cell functional heterogeneity[5] observed in patients. This already offers critical advantages for pre-clinal modelling and uncovering leads to improve CAR T cell treatment[3,88] one of the most promising therapeutic avenues[105] for a disease that has not seen any clinical improvement in decades[2] .

#### Acknowledgements

All imaging was performed at the Princess Máxima Imaging Center. We thank the Princess Máxima Center Single Cell Genomics Facility, as well as the Leiden Genome Technology Center, for performing sc/snRNA-sequencing, R. Moeniralam and E. de Boed from the Princess Máxima Center Pathology Diagnostic Laboratory for performing immunohistochemical staining and the flow cytometry facilities at the Princess Máxima Center and Laboratory of Translational Immunology (UMCU) and Mara Nicolasen for cell sorting. We thank J. Lammers for providing gene expression data from patient material, Z. Odé for her collaboration on establishing cerebral organoids, H.R. Johnson and A. Zomer for assistance with *in vivo* experiments and J. Bunt for sharing his knowledge on neural development. This work was financially supported by the Princess Máxima Center for Pediatric Oncology and Oncode Institute, the Netherlands. A.C.R was supported by an ERC-starting grant 2018 project (no. 804412) and N.B. by a research grant from Stichting Proefdiervij.

#### Competing interests

A.C.R. and N.B. are listed as inventors on a pending patent related to the novel brainstem-regionalized organoid model. A.K.L.W., E.J.W., M.A. and A.C.R. are listed as inventors on a pending patent related to the development of marker-based T cell selection.

#### Author Contributions

N.B. and A.C.R conceptualized the work with critical input from H.C.R.A, A.K.L.W., N.D., C.R.M., H.S., C.H., M.A., and A.A., and wrote the manuscript with support from E.J.W..

H.C.R.A., C.H. and N.B. grew pontine organoids and performed DMG tumour induction. H.C.R.A., C.R.M., N.B., S.d.B. and M.A. analysed bulk and scRNA-seq datasets with critical input from H.S. and F.F.K.. N.D., H.C.R.A. and N.B. performed and analysed lineage tracing and cNMF experiments supervised by A.A. and A.C.R.. C.M. provided TrackerSeq and critical input for analysis of lineage tracing experiments. N.B., H.C.R.A., and M.B.R. performed multi-spectral 3D imaging, using specific technology provided by A.E.. T.B. performed cyclic immunofluorescence imaging, R.V.U.C. live cell imaging, D.J.K. flow cytometry and S.P. and E.M. DNA methylation profiling. A.K.L.W. and C.H. integrated microglia and performed all GD2 CAR T cell experiments with help from A.M.C. for T cell expansion and L.C.D.E. for cell sorting. A.K. provided GD2 CAR T cells and J.K., Z.S. and S.N. protocols for T cell rapid expansion. E.J.W., A.K.L.W., F.F.K., E.v.V. and M.A. analysed DMGO CAR T cell treatment data, including single scRNA-seq analysis of GD2 CAR T cells and microglia. M.E.G.K. contributed critical knowledge on DMG histopathological features and performed immunohistochemical staining on DMGOs. M.R. and M.K. provided healthy brain organoids as a refence dataset for iCNV analysis and knowledge related to brain organoid generation. S.d.B. offered critical computational support and infrastructure for data analysis.

#### Methods

##### Ethics

For the use of all DMG patient samples, patients and/or parents or guardians provided written informed consent according to national laws and in agreement with the declaration of Helsinki (2013). This study is Institutional Review Board (IVB) approved and registered under national registry number 2020.142.

##### Stem cell culture

Brain organoids were generated from 3 different cell lines encompassing human Embryonic Stem Cells (hESCs) H9 (WA09, WiCell) and H1 (WA01, WiCell) (both derived from human blastocysts[106]) and induced Pluripotent Stem Cells (iPSCs) C7-a (RUID 06C52463, derived from $CD4^{+}$ T cells). The iPSC line C7-a was obtained from Rutgers University Cell and DNA Repository (RUCDR) and contained a Cre-inducible H3.3K27M reading frame in the endogenous *H3F3A* locus[19]. Cell lines were cultured in mTeSR Plus medium (Stem Cell Technologies, Cat. #100-0276) and incubated at 37°C with 5% $CO_2$. The cells were grown on Matrigel-coated (Corning, Cat. #354277) 6-well plates and passaged when 70-80% confluent by non-enzymatic detachment of colonies using Gentle Cell Dissociation Reagent (GCDR, Stem Cell Technologies, Cat. #100-0485). All cell cultures were routinely tested for the presence of Mycoplasma species.

**Embryoid body formation**

The different stem cell sources were washed with 1X Dubecco's Phosphate-Buffered Saline (DPBS, Gibco, Cat. #14190144) and detached with GCDR, before spinning down at 300 RCF for 5 minutes. Next, the cells were resuspended in BASE medium (1:1 Advanced DMEM/F-12 medium (Gibco, Cat. #12634010) and Neurobasal medium (Gibco, Cat. #10888022), 1X GlutaMax (Gibco, Cat. #35050061)) and counted. 70,000 cells/ml were added to DAY 0 medium (BASE medium, 10 µM Y-27632 (ROCKi, AbMole BioScience, Cat. #M1817), 4 ng/ml Fibroblast Growth Factor 2 (FGF2, PeproTech, Cat. #100-18C)). For Embryoid Body (EB) formation, 7,000 cells in 100 µl medium are seeded per well of an ultra-low attachment (ULA) treated U-bottom 96-well plate (Nexcelom, Cat. #ULA96U020/PHC Europe B.V., #MS-9096UZ) and incubated at 37°C with 5% $CO_2$. From day 2 – day 21, PATTERNING medium (BASE medium, 1X N2 (Gibco, Cat. #17502048), 1 mg/ml Heparin Solution (Stem Cell Technologies, Cat. #07980) was used.

**Organoid patterning**

To induce brainstem identity, organoids were patterned using timely additions and replacement of morphogen supplemented media. In week 1, WEEK 1 medium (PATTERNING medium, 50 ng/ml FGF2, 1 µM Dorsomorphin (DM, Stem Cell Technologies, Cat. #72102), 10 µM SB431542 (SB43, Stem Cell Technologies, Cat. #72232), 3 µM CHIR99021 (CHIR, Stem Cell Technologies, Cat. #72052)) was used. On day 2, 100 µl WEEK 1 medium was added per well. On day 5, 100 µl medium per well was replaced with fresh WEEK 1 medium. In the second week, WEEK 2 medium (PATTERNING medium, 1 µM DM, 10 µM SB43, 3 µM CHIR, 10 ng/ml Fibroblast Growth Factor 4 (FGF4, Stem Cell Technologies, Cat. #78103.1), 10 µM All-Trans Retinoic Acid (RA,

Stem Cell Technologies, Cat. #72262), 1 µM Purmorphamine (PMA, Stem Cell Technologies, Cat. #72202) was used. On day 7, 190 µl medium was replaced with fresh WEEK 2 medium and on day 9 100 µl medium was replaced. On day 11, the EBs were embedded in 12 µl Matrigel droplets and 5 droplets were transferred to each well of a 12-well suspension plate (Greiner Bio-One, Cat. #665102) with 1 ml WEEK 2 medium and incubated at 37°C with 5% $CO_2$. In week 3, WEEK 3 medium (PATTERNING medium, 10 ng/ml FGF4, 10 µM RA, and 1 µM PMA) was used. Until day 21, every 2 days, the medium was refreshed with WEEK 3 medium. On day 17, the plates were placed on an orbital shaker inside a 5% CO2 incubator at 37°C. From day 21 onwards, every 2-3 days, the medium was refreshed with MATURATION medium (1:1 Advanced DMEM/F-12 medium and Neurobasal medium, 1X GlutaMax, 0.5X N-2, 0.5X B27 without vitamin A (Gibco, Cat. #12587010), and 1X Penicillin-Streptomycin (Pen-Strep, Gibco, Cat. #15140122)).

**DMG driver mutation-expressing and genetic lineage tracing plasmids**

To induce DMG tumour growth in hESC-derived pontine organoids, the following plasmids were used: pCAGPbase, PBCAG_DNp53_IRES_luciferase, PBCAG_PDGFRA-D842V_IRES_eGFP, and PBCAG_H3K27M_eGFP. Alternatively, to induce DMG tumour growth in iPSC-derived pontine organoids, the H3K27M-expressing plasmid was replaced with 1.00 µg/µl Ssi-Cre to induce with an inducible H3.3-K27M mutation targeted to the endogenous histone locus. As a control, the following plasmids were used: 1.50 µg/µl pCAGPbase and 1.50 µg/µl PB_Venus. All plasmids were kindly provided by the Pheonix laboratory[16]. For genetic lineage tracing, 1.50 µg/µl TrackerSeq[107] was added to the tumour and control plasmid mix.

##### ***In situ* electroporation and monitoring of tumour growth**

On day 11, unless stated otherwise, brainstem organoids were injected with a mixture of plasmid DNA (1.50 µg/µl per plasmid) and 0.1% (w/v) FastaGreen (Merck, Cat. #F7252-5G) using a FemtoJet 4i (Eppendorf, Cat. #5252000013) with the following parameters: Injection pressure (Pi) = 15 hPa and compensation pressure (Pc) = 5 hPa. Subsequently, the organoids were electroporated using a NEPA21 Super Electroporator (Nepagene) and CUY650P1 (Nepagene) tweezers with the following parameters: Voltage = 50 V, pulse length = 10 ms, pulse interval = 50 ms, number of pulses = 4, and decay rate = 10%. Transfer Pulse; Voltage = 20 V, pulse length = 50 ms, pulse interval = 50 ms, number of pulses = 5 and decay rate = 40%. Using the impedance (kΩ) measurement of the NEPA21 Super Electroporator, voltage was automatically re-adjusted to optimize cell perforation and viability per individual organoid. Electroporation was performed by applying a shock twice in orthogonal direction. After electroporation, the organoids were incubated at 37°C with 5% $CO_2$ for at least two hours to recover before Matrigel embedding. To monitor tumour growth over time, organoids were imaged on a Leica DM IL LED microscope with an N PLAN 5x/0,12 PH0 objective and compared to the mVenus-positive control.

##### **DMGO orthotopic transplantation**

All murine experiments were conducted in compliance with the Animal Welfare Committee of the Princess Máxima Center for Pediatric Oncology based on local and international regulations. 3–4-week-old NSG mice were anaesthetized using Isolurane/$O_2$ inhalation and transferred to a stereotaxic frame. Eye ointment was applied, and 0.05 mg/kg Buprenorphine injected subcutaneously. After removing hair from the surgical site, a 1 cm incision was made in the skin to expose the skull and 3 mg/kg Lidocaine was applied topically. Under a stereo microscope, a Dremel was used to drill a circular groove of 5 mm in the skull above the right cerebral cortex. Cortex buffer was applied before dura mater and 2 mm^3 brain tissue was removed to accommodate the DMGO transplant. DMGOs were pre-selected based on GFP signal 1-2 weeks after electroporation and, if too big in size, cut in half before transplantation. After placing the DMGO, the brain was covered with a neuro-patch, the skull closed with dental cement, and the wound closed using skin glue. After surgery, 0.06 mg/ml Carprofen was provided in the drinking water for 3-5 days and mice were monitored 2-3 times per week for signs of weight loss, lack of grooming, and/or reduced mobility. If mice reached study- (21 days) or human endpoint based on the monitoring of symptoms, they were put under deep anaesthesia by intraperitoneal injection of a 75 mg/kg Ketamine + 1 mg/kg Medetomidine mix solution. Trans-cardiac perfusion was performed with PBS and 4% PFA and after resection brains were cut into 300 um sections using a vibratome. Staining, clearing, and imaging was performed as described below with the following specific primary and secondary antibodies: CD31 (Abcam, #ab134168), Anti-Rabbit AF488 (ThermoFisher, #A21206), GFP booster ATTO647N (ChromoTek, #GBA647N-100).

**Multi-spectral large-scale single-cell resolution 3D (mLSR-3D) imaging**

A comprehensive list of buffers, products, and clearing agents used for sample preparation can be found in the protocol by van Ineveld *et al.*[108]. In short, organoids were fixed in 4% paraformaldehyde (PFA, Sigma-Aldrich, Cat. #441244) for 30 minutes at 4°C, washed 3X in PBT (1:1000 Tween-20 in 1X PBS) for 15 minutes at 4°C, embedded in 4% low melting point (LMP) agarose (Invitrogen, Cat. #16520-050), and sliced into 100-250 µm sections using a Leica VT 1200

S Vibratome. Sliced organoids were permeabilized in washing buffer 1 (WB1) on a shaker at 4°C for 3 hours and, subsequently, stained with primary antibodies diluted in washing buffer 2 (WB2) on a shaker overnight at 4°C. After primary antibody staining, the slices were washed with WB2 at 4°C for 5 hours and stained with secondary antibodies diluted in WB2. All combinations of primary and secondary antibodies used are listed in **Supplementary Table S7**. Additionally, the cell nuclei and membranes were stained with DAPI (1:2000, Invitrogen, Cat. #D1306) and Phalloidin-Atto 488 (1:400, Invitrogen, Cat. #A12379), respectively. After secondary antibody staining, the organoid slices were washed with WB2 for 5 hours at 4°C and cleared with the fructose-based clearing agent FUnGI according to the protocol by Rios *et al.*[109] and mounted on coverslips with silicone as spacer. The slices were imaged using a Zeiss LSM 880 Confocal microscope with a 25X (NA 0.8) objective and Leica Stellaris with 20X (NA 0.75) and 40X (NA 1.3) objectives. Alternatively, intact organoids were fixed and cleared using the organic solvent-based vDISCO method according to the protocol by Cai *et al.*[110] and imaged using a Leica SP8 microscope with a 16X (NA 0.6) BABB-compatible objective.

**2D FFPE imaging**

DMG patient material, as well as DMGOs, were fixed in formalin and embedded in paraffin at the histopathology department of Princess Máxima Center for pediatric ncology to obtain formalin-fixed paraffin-embedded (FFPE) tissue for WHO-standardized tumour classification. Patient and organoid material was sliced into 3 µm sections prior to hematoxylin and eosin (H&E) and subsequent stainings. Immunohistochemical staining was performed on the Leica BOND RX Fully Automated Research Stainer using the Bond Polymere Refine Detection kit (Leica, Cat. #DS9800). The following antibodies were used: GFAP (RTU, Leica, Cat. #PA0026), NF (RTU, Leica, Cat.

#PA0371), H3K27M (1:400, Abcam, Cat. #AB190631), and H3K27me3 (1:200, Cell Signaling, Cat. #9733S). Stained tissue sections were analysed by an experienced neuropathologist. Sections for fluorescent-labelled imaging were subjected to antigen retrieval using Target Retrieval Solution pH 9 (Agilent Dako, Cat. #S2367) with 60 minutes boiling time and stained using the mLSR-3D protocol with reduced incubation times (1h incubation at RT). Combinations of primary and secondary antibodies used are listed **in Supplementary Table S7**. Additionally, cell nuclei were stained with DAPI (1:2000) and the slices were imaged using a Leica Stellaris confocal microscope with a 20X (NA 0.75) and 40X (NA 1.3) objective.

#### DNA methylation profiling

The DNA methylation profile of a pooled DMGO sample consisting of 3 independent replicates was compared to cases of DMG, glioblastoma and posterior fossa ependymoma obtained from published datasets[111,112]. Data was loaded in R environment (v4.3.1), probe filtering performed using package ChAMP[113] and each array platform was processed separately (HumanMethylation450, or EPIC) using method "minfi"[114] and filtering out probes located on SNPs, sex chromosomes or with detection p-value > 0.01. Raw beta values were merged using function combineArrays and normalized with method BMIQ[115]. We selected the 10,000 probes with the highest standard deviation and calculated the Pearson correlation between samples, weighted by the inverse of variance. This resulting correlation matrix was used to compute a distance matrix, which served as the input for the Rtsne function from the Rtsne package.

#### Cyclic immunofluorescence imaging

FFPE tissue sections of organoids were deparaffinized in Xylene (3 x 3 min) followed by rehydration in a series of graded alcohol for 1 min each (2 x 100%, 2 x 95%, 1x 70%). Sections were washed in deionized water (2 x 1 min) and put in Target Retrieval Solution, pH 9 (Agilent Dako). Antigen retrieval was performed for 40 min at 95°C. The sections were allowed to cool down to room temperature and were washed for 5 min in deionized water followed by storage in PBS until further use. Cyclical immunofluorescence imaging was performed as previously described[116] . After antigen retrieval, a barrier was drawn surrounding the tissue using a hydrophobic pen. Tissue was exposed to a blocking solution consisting of 100 mM $NH_4Cl$ (ThermoFisher), 150 mM Maleimide (Merck), and 10% donkey serum (Merck) in PBS for 1 h in a humidified chamber at RT. Next, the blocking solution was replaced with a primary antibody staining solution containing 100 mM $NH_4Cl$ and 5% donkey serum. Primary antibodies used were IBA1 (Fujifilm Wako, 019-19741), CD163 (ThermoFisher, MA5-11458), SPP1 (R&D Systems, AF1433-SP), and P2RY12 (Atlas, HPA014518). Sections were incubated for 2.5 h in a humidified chamber on an orbital shaker followed by washing in PBS (3 x 5 min). Sections were exposed to a secondary antibody staining solution (containing 100 mM $NH_4Cl$, 5% donkey serum, and DAPI (Biolegend)) for 1 h at RT in the dark. Secondary antibodies used were Anti-Rabbit Cy5 (Jackson ImmunoResearch, 711-175-152), Anti-Goat AF555 (ThermoFisher, A32816), Anti-Mouse AF488 (ThermoFisher, A21202). After washing (3 x 5 min) in PBS, SlowFade Gold antifade mounting medium (Invitrogen) was applied on the sections. Imaging was performed on a Leica DMi8 Thunder imaging system with a HC PL APO 20x/0.80 objective. After imaging, the coverslips were removed in PBS followed by washing in PBS (3 x 5 min). Antibody removal was performed by applying elution buffer (Lunaphore) to the sections for 3 min. After washing in PBS (3 x 5 min), the following imaging cycle was started by again applying the blocking buffer.

Images of each cycle were aligned based on the DAPI signal using a previously developed tool, available at https://github.com/Dream3DLab/CycFluoCoreg. In brief, the tool applies affine transformation to each channel to align images to the DAPI reference followed by B-spline transformation, refining the alignment. The resulting composite images were imported into QuPath (v0.4.4)[117] where nuclei were detected and segmented using a cell expansion of 2.5 μm. An object classifier using RandomTrees was trained for each marker on two separate images. These object classifiers were combined into a composite classifier that was applied to all images. The resulting dataset containing the count of classified cells in each image was exported to R for quantification and visualization.

**Bulk RNA sequencing**

Brianstem organoids were used for bulk RNA sequencing at different patterning and maturation time points (week 1, 2, 3, 4, 8 and 12) from at least 3 different batches and generated from different stem cell sources, either H9 or C7-a. Bulk RNA sequencing was performed on pooled organoids, which were collected in 2 ml DNA low-binding tubes (Eppendorf, Cat. #0030108078). In addition, brainstem organoids that were patterned with low (10 ng/ml) and higher (20 ng/ml) concentrations of FGF2 or FGF4 from week 2 to 3 were collected at the end of week 3. Organoids were mechanically dissociated in Hank's Balanced Salt Solution (HBSS, Gibco, Cat. #14025092), before spinning down at 800 RCF for 5 minutes at 4°C. Subsequently, supernatant was removed, and the cell pellet snap-frozen on dry ice and stored at -80°C. Total RNA was extracted from cell pellets using the RNeasy Mini Kit (Qiagen, Cat. #74104) according to manufacturer's instructions. RNA concentration and integrity were evaluated by running 1 μl of the total RNA sample on an RNA 6000 pico gel (Agilent, Cat. #5067-1513) using a 2100 Bioanalyzer (Agilent, Cat.

#G2939BA). Samples with an RNA Integrity Number (RIN) above 8 were sequenced on the Illumina NextSeq500 platform and subsequently mapped and aligned by the Utrecht Sequencing Facility (Useq).

**Bulk RNA data analysis**

Raw counts of bulk sequencing data were combined into a single matrix and normalized by DESeq2 median-of-ratios and variance-stabilized transformation (vst) (v1.40.2)[118]. Gene counts below 10 were filtered out. To determine organoid batch variability, Spearman's Rank Coefficient was computed using built-in R functions (package "stats"). For principal component analysis (PCA) single organoid samples of the same batch were merged. Barplots were generated using top 15 genes per region on custom level 2 and 3, while spatial maps were generated using top 15 genes on custom level 1.

**Single-cell RNA sequencing sample preparation**

Brainstem organoids from the same batch were dissociated at selected timepoints to capture various stages of development. Single-cell suspensions were collected from the following stages: embryoid body (day 5), neural induction (day 11), post-Matrigel embedding (day 14), and neuronal specification/maturation (days 20, 30, 60, 90, and 120). To ensure sufficient cell numbers, multiple organoids were pooled, with 24 organoids used for earlier timepoints and as few as 7 for later stages. For timepoints after Matrigel embedding, Cell Recovery Solution (Corning, 354253) was used to dissolve the Matrigel. Organoids were incubated in this solution at 4°C for 15 minutes, halved, and washed in HBSS without $Ca^{2+}$ and $Mg^{2+}$. The dissociation process utilized the Neural

Tissue Dissociation Kit (Miltenyi Biotec, 130-092-628), which is papain-based. Briefly, pre-warmed papain buffer was added to the organoids and incubated at 37°C for 15 minutes in a rocking incubator. Enzyme Mix A was then added, and the suspension was triturated 15 times using wide-bore and P1000 pipette tips. The mixture was incubated further with regular visual inspection for approximately 10 minutes, or until a single-cell suspension was achieved. After dissociation, cells were filtered through 70 µm and 20 µm pre-separation filters to remove debris. The filtrate was centrifuged to pellet the cells, which were then washed by resuspension in HBSS without $Ca^{2+}$ and $Mg^{2+}$. Cell counts were performed using a Trypan Blue assay on the automated Countess Cell Counter (Thermo Fisher Scientific). For storage, the cell suspension was divided into two aliquots. After pelleting, the cells were resuspended in 1 mL of mFreSR cryopreservation medium and frozen at -80°C for 24 hours. Subsequently, cryotubes were transferred to liquid nitrogen for long-term storage until scRNA seq was conducted.

**Library preparation and single-cell RNA sequencing**

On the day of sequencing samples were thawed by warming the cryovials in a 37°C water bath until only a small clump of ice remained. The contents were then transferred to 10 mL of pre-warmed DMEM containing 10% FBS and centrifuged to pellet the cells. The cells were washed twice with PBS containing 5% BSA and filtered through a 40 µm Flowmi cell strainer to remove debris and aggregates. Cell viability and counts were assessed using a Trypan Blue assay on the automated Countess Cell Counter (Thermo Fisher Scientific). After counting, cells were resuspended in an appropriate volume to target the capture of 30,000 cells. Single-cell RNA sequencing libraries were generated using the Chromium Single Cell 3' v4 Library & Gel Bead Kit (10x Genomics) following the manufacturer's protocol and sequenced on the Illumina

NovaSeq platform. A summary of all single-cell sequencing experiments is provided in **Supplementary Table S1**.

**Count matrix generation and preprocessing**

Transcript count matrices were generated using Cell Ranger (v7.0.1) with default parameters, aligning the sequenced reads to the 10X Genomics provided reference genome (hg38). Count matrices were further processed using the Seurat R package. To ensure data quality, cells were filtered based on mitochondrial content, the number of detected genes, and the number of unique molecular identifiers (UMIs). Thresholds for filtering were calculated per sample as follows:

- The minimum number of detected genes was set as the greater of 400 or two standard deviations below the mean.
- The maximum number of detected genes was set as two standard deviations above the mean.
- Mitochondrial content was capped at the greater of 5% or two standard deviations above the mean.
- The minimum UMI count was set as the greater of 0 or two standard deviations below the mean, while the maximum was set as two standard deviations above the mean.

These thresholds were computed dynamically for each sample to account for variations in dataset quality, and specific values for each sample are provided in **Supplementary Table S2**. Cells passing all filters were retained for further analysis. Filtered transcript counts were normalized to the total number of counts per cell, scaled to 10,000 UMIs per cell and log transformed.

#### Benchmarking integration

To establish a multi-level initial annotation for label-aware integration, we utilized Snapseed alongside a predefined set of marker genes[40]. Annotation was performed using the annotate_hierarchy() command with default settings. Employing 3,000 highly variable genes (HVGs) and timepoint as a batch covariate, we used the scib-metrics package to evaluate integration performance across several methods, leveraging GPU-acceleration when applicable. For semi-supervised integration approaches the coarse level pre-annotion derived from snapseed was used as initial guidance. The following representations were compared for integration performance:

- Unintegrated PCA
- Harmony (as implemented in Scanpy)
- Harmony-Timeseries (as implemented in Scanpy)
- Batch Balanced KNN (BBKNN, neighbors_within_batch=3)
- Scanorama
- Mutual Nearest Neighbors (MNN) (as implemented in Seurat)
- scVI (batch_size=1024, max_epochs=500)
- scANVI (using Snapseed level 1, max_epochs=100)
- scPoli (using Snapseed level 1, n_epochs=50, pretraining_epochs=40)

scPoli was selected on its weighed performance in batch correction and preserving biological variance. Consequently, the scPoli-derived latent space was selected for downstream analysis. For visualization, PAGA was used to generate a coarse-grained graph representation of the data. This graph provided and overview of the dataset's global structure and informed the construction of the final UMAP embedding. The UMAP served as the primary visualization for interpreting the integrated dataset.

**Reference mapping against the HNOCA**

The brainstem organoid integrated time course dataset was aligned to the 3,000 highly variable genes (HVGs) of the human neural organoid cell atlas (HNOCA)[40] . For genes not present in the brainstem dataset, expression values were filled with zeros. The original scPoli model was downloaded from the publication's GitHub repository (https://github.com/theislab/neural_organoid_atlas/tree/main/supplemental_files/scpoli_model_params) and loaded into the mapper module of the HNOCA-tools package, v0.1.1. The organoid dataset was projected into the HNOCA space using the map_query() method with the following parameters: retrain='partial', batch_size=256, unlabeled_prototype_training=False, n_epochs=100, pretraining_epochs=90, eta=10, alpha_epoch_anneal=10. After projection, a weighted k-nearest neighbor (wkNN) graph was computed with $k$ = 100. This neighbor graph facilitated the transfer of cell labels from the HNOCA dataset and the generation of a shared UMAP representation that combined the HNOCA and brainstem organoids datasets. Finally, presence scores were calculated using the get_presence_scores() function from the HNOCA-tools package, providing insights into the representation of brainstem organoid cells within the HNOCA framework.

**Reference mapping against the HDBCA**

The CellRanger-processed dataset of the human developing brain cell atlas (HDBCA)[41,42] was downloaded via the link provided on the publication's GitHub page. Cells with fewer than 300 detected genes were excluded from further analysis. The dataset was normalized by total counts per cell, scaled, and log-transformed. The HDBCA gene expression space was then intersected with the gene set from the HNOCA. A higher-resolution, cluster-based cell type annotation of the HDBCA, recently published[42] , was integrated into the HDBCA dataset. A scVI model was trained on the 2,000 HVGs of the HDBCA with 'donor_id' specified as the batch key. The following hyperparameters were used: n_latent=20, n_layers=2, n_hidden=256, use_layer_norm='both', use_batch_norm='none', encode_covariates=True, dropout_rate=0.2. The model was trained with a batch_size=1,024 and early_stopping=True for a maximum of 500 epochs or until convergence. The trained scVI model was subsequently fine-tuned with scANVI, using 'CellClass_Mossi' as cell type labels. Fine-tuning was conducted for 100 epochs with batch_size=1,024, early_stopping=True, and n_samples_per_label=100.

To compare the brainstem organoid model with the HDBCA, the brainstem organoid dataset was aligned to the 2,000 HVGs used to train the scANVI model. Missing gene expressions in the brainstem organoid dataset were imputed with zeros. The aligned brainstem organoid dataset was loaded into the trained scANVI model, and a query model was trained with the following parameters: batch_size=1,024, max_epochs=100, weight_decay=0.0. Cell type labels were then transferred to the brainstem organoid dataset using the scvi-tools predict() method.

**HNOCA comparative abundance analysis**

The scCODA algorithm, as implemented in the pertpy package v0.9.4, was employed to analyze cell type compositional changes between the brainstem organoid model and HNOCA datasets. Timepoints prior to day 30 were excluded as these include mostly progenitors, along with pluripotent stem cells, neuroepithelial cells, and non-neural lineages. For the analysis, the 'publication_id' key was used as the covariate_obs, and 'bio_sample' was specified as the sample_identifier. Cell type annotations were grouped by region. For cell types lacking a regional annotation, the cell type transferred from the HNOCA dataset was used. scCODA was executed iteratively, with each cell type selected once as the reference, using default parameters and the No-U-Turn Sampler (NUTS) for Bayesian inference. A majority voting system identified cell types that were credibly differentially abundant, defined as those deemed significant in more than half of the iterations.

**Gained presence analysis**

To validate the regional identity of the brainstem organoid model in the context of the HNOCA, presence scores were calculated for both datasets using the HDBCA as a reference, following the method described in He *et al.*[40] . The presence score quantifies how often a cell type in the HDBCA is observed in either the brainstem organoid model or the HNOCA dataset. For the brainstem organoid model, the presence score was computed by summing the weights of each reference cell linked to the query cell in the wKNN graph. These raw scores were smoothed using a random-walk-with-restart procedure and subsequently log-transformed. Scores below the 5th percentile or above the 95th percentile were clipped, and the clipped values were normalized to a range of [0,1]. Presence scores for the HNOCA were calculated as described in He *et al.*[40] . The gained presence score was determined by subtracting the HNOCA presence scores from the brainstem organoid presence scores for each cell in the HDBCA. Only positive differences, reflecting an increased presence of HDBCA cell types in the brainstem organoid model relative to the HNOCA, were considered. These gained presence scores were averaged across each HDBCA cluster and grouped by region for visualization.

**Glial differential expression analysis**

Differential expression (DE) analysis was done to evaluate and compare the transcriptomic similarity of the glial lineages in our brainstem organoid model. Cells expressing more than 200 genes and belonging to either OPC or Glioblast lineages in the HDBCA, HNOCA and the brainstem organoid model were aggregated into a pseudobulk object with three pseudoreplicates per dataset using the python implementation of decoupler. egdeR was used to compute DE genes for each glial lineage using HDBCA_vs_HNOCA and HDBCA_vs_brainstem organoids respectively, correcting for cell numbers, as well as median and standard deviation of the number of detected genes per pseudobulk sample. Genes were tested using the Genewise Negative Binomial Generalized Linear Model implemented in edgeR. Genes with an absolute log2FC above 1 and a qvalue below 0.05 were labeled as DE. For DE gene distribution between the HNOCA and the brainstem organoid model, scipy's fisher's exact test was used to calculate the odds ratio and p-value.

**Regional identity analysis using Voxhunt**

To compare scRNA seq data from the brainstem organoid model with mouse spatial gene expression data, the VoxHunt R package was utilized. Spatial gene expression samples from embryonic day (E) 13 and E18 were selected for the analysis. For each region at annotation level 'custom_1', the top 10 genes were identified based on their provided Area Under the Curve.

**Single cell RNA sequencing of tumour and GD2 CAR T cells and Trackerseq libraries**

We performed single cell dissociation using the Neural Tissue Dissociation Kit (P) (Miltenyi Biotec, Cat. #130-092-628), adjusted for DMGOs. After cutting the organoids into smaller pieces and adding the enzyme mixes, the samples were incubated on an orbital shaker at 37°C and resuspended in regular intervals with a P1000 until a single cell suspension was reached, which was verified under the microscope using trypan blue to check for cell death. After dissociation, the single cell suspension was washed twice with PBS +/+ (magnesium/calcium+ 3% FBS). Next, DAPI was used as a viability dye (DAPI 1:5000). Stained single cells suspensions were filtered using a 40 µm Flowmi cell strainer and sorted by FACS on DAPI exclusion and enriched for GFP expression for tumour cells. For T cells, see section 'FACS of GD2 CAR T cell treated DMGOs'. FACS experiments were performed using the CytoFLEX SRT Benchtop Cell Sorter (Beckman Coulter).

All samples - analysed simultaneously using Hash tag oligo (HTO) demultiplexing (described in subsequent section) - were pooled together, and single-cell encapsulation was performed according to the manufacturer's protocol 10X Genomics (Cell Preparation for Single Cell, Demonstrated Protocol, CG000053). Pooled cDNA amplification was generated using the Chromium Single Cell 3' V3 Library & Gel Bead Kit using the TruSeqR1, TruSeqR2 and partial TSO (template switch oligo) standard primers. Total cDNA of this reaction was then used for Library preparation for mRNA and hashtag oligos as described per the manufacturer's protocol. For the lineage tracing strategy, single-cell gene-expression, hashing, and TrackerSeq lineage barcode 10x genomics barcoded libraries were constructed for 60-day post-electroporation DMGOs. nested PCRs were additionally used to further amplify the lineage barcodes (primers in first PRC (10 cycles): sequences: 5'-CTACACGACGCTCTTCCGATCT-3 (Read1-Forward primers from 10X, CTTCTCGTTGGGGTCTTT (eGFP primer-Reversed) Annealing Temperature was 60 and elongation time 30 sec; primers in the second PCR (10 cycles) Annealing Temperature was 60 and elongation time 30 sec: standard P5-Read 1_Forward and Fun series 70x primer-Reversed from 10x). Sequencing of the prepared libraries was performed on an Illumina NovaSeq6000 in PE150 mode, and raw fastq files were processed and mapped with CellRanger v3.1.0. using a custom reference hg38 genome including sequences used in the electroporation methods, namely EGFP, H3.3K27M, Dnp53, Luciferase and Pdgfra.D824V.

**Hash tag oligo (HTO) demultiplexing**

Each organoid was stained with a different TotalSeq-A anti-human hashtag (Hashtag A0251-A0265, Biolegend), allowing to tag each cell with a sample-specific artificial oligonucleotide that can be recovered by sequencing. The hashing oligos were designed to recognize most human cells using a combination of two clones against CD298 and β2 microglobulin. In sum, the cell pellet was resuspended in staining buffer (50 ul for 500,000 cells). Unspecific binding was reduced by adding 5ul of human Fc blocking reagent to the sample (FcX Human truStain, Biolegend Cat. #422301). After 10 min incubation at 4 °C, 1 μl of a unique cell hashing antibody was added to each sample and incubated for 20 min at 4 °C and then washed 3x using PBS +0.04% of BSA. Demultiplexing of HTO-hashed single cell RNA sequenced samples was done using Seurat (v4.4.0). Briefly, CellRanger derived count matrices for `Gene Expression` and `Antibody

Capture` were loaded and the Unique Molecular Identifiers (UMI) intersected to filter cells that are detected in both matrices. The `Gene Expression` count matrix was used to generate a Seurat object and the `Antibody Capture` was added as an assay ("HTO"). The "HTO" assay was normalized using Centered Log-Ratio (CLR). The Seurat object was subsequently demultiplexed with the embedded HTODemux() function using default parameters. Cells which were assigned one HTO-barcode (Singlets) were subsetted and used for further analysis.

**Pre-processing and curation of single cell datasets**

Downstream processing was done on individual samples using the Seurat workflow. Cells were filtered on mitochondrial content (indicative of dying cells), number of detected genes and number of UMIs. Threshold for these parameters were defined per sample based on their distribution. One sample with less than 100 cells after filtering was excluded from subsequent analysis. A gender score was assigned to each cell based on the expression of XIST using the AddModuleScore() function in Seurat. Cells after filtering were scaled to 10.000 UMIs per cell and log-normalized. Mitochondrial content, gender and UMI counts were regressed out from normalized gene counts, the genes scaled and centred. Dimensionality reduction was applied to the top 2,000 highly variable genes using PCA. The first 30 principal-components were used for projection in Uniform Manifold Approximation and Projection (UMAP) space, for construction of a shared nearest neighbour (SNN) graph and clustering based on the Louvain algorithm. Potential doublets were detected using DoubletFinder3 (v2.0.3) and excluded. Cell cycle phase was determined as implemented in Seurat. Samples were visually inspected for expression of cell type marker genes and expression of tumorigenic plasmid genes. One sample contained high mesodermal markers

(*PAX7*, *MYOG*, *MYOD1*) and was therefore removed from the dataset. The remaining cells were used for integration and analysis.

**Integration of single cell data**

After pre-processing, all datasets were merged into one single Seurat object. The top 2,000 highly variable genes were recalculated, and the Seurat object subsetted to only contain these genes. Mitchondrial and ribosomal content, gender, cell cycle phase and UMI counts were regressed out from the merged normalized counts and the genes scaled and centred. The Python package scVI (v1.0.4) was employed using reticulate (v1.34.0) to balance confounding factors driving differences between the multiple datasets. To do this, the Seurat object was converted to AnnData format (sceasy, v0.0.7) and loaded into a scVI model with "RunID" as batch variable. The model was trained for 400 epochs and after training the 10 latent embeddings were extracted and added to the Seurat object as a DimReducObject. These embeddings were used to project the cells in UMAP space using default parameters. A SNN graph was constructed, and clusters detected using the Louvain algorithm on different resolutions. Clusters were inspected by building a clustertree using clustertree (v0.5.1) and representation in UMAP space.

**Inferred copy number variation (iCNV) analysis**

To confirm tumorigenicity, we inferred the copy number of variation (iCNV) from single-cell gene expression data using SCEVAN (v 1.0.1). SCEVAN utilizes a sliding window approach across the genome, comparing the expression levels in test cells to reference cells. On a sample-by-sample basis, we used a non-transfected organoid as a healthy reference to estimate iCNVs in the malignant cells. The iCNV result is displayed in a heatmap representing CNV inference across the genome. Elevated or reduced expression levels indicative of CNVs are visually represented, facilitating easy identification of malignant clusters.

#### Annotation and reference comparisons

Samples from different sequencing runs were integrated using the scanpy implementation of harmony. On the integrated representation a diffusion map was calculated which served as input for a ForceAtlas representation of the data. Leiden clustering using resolution=1 revealed 16 clusters, which were annotated using tumour state marker genes derived from Liu et al[13] . Annotation was subsequently confirmed by projecting the tumour into two different patient datasets[12,13] using Azimuth. Briefly, raw expression matrices from reference datasets were obtained and processed using Seurat, as described in the corresponding papers. Tumour data was projected onto these reference datasets using the RunAzimuth function from the Azimuth package (v.0.5.0).

For comparison of the DMGO tumour model to other tumours or models[11–13,52] multiple references were merged, normalized and scaled. PCA was obtained using RunPCA() and annotation was unified and condensed. Anchors were obtained by FindTransferAnchors(), using the first 30 PCs of the shared PCA space and all genes of the reference dataset as features, and used to transfer the reference annotation to the query tumour data. The labels were transferred to the query Seurat as a separate assay and used as input for visualization.

#### Gene set enrichment analysis

The AddModuleScore() function in the Seurat package was used with default settings to compute gene set enrichments from curated lists of marker genes which were subsequently visualized. To compare gene expression between DMGO and reference datasets, the raw counts from the objects were merged, normalized, and scaled as one object before visualizing the gene of interest. All sequencing analysis was performed on R (v4.3.1) in RStudio (2023.090+463) or Python (v3.10.4). Molecular signatures for spatially restricted oligodendrocyte precursor lineages retrieved from Braun *et al.*[41] (**Supplementary Table S5c**) were used for similarity comparison of cNMF generated Program 1 and 2.

**TrackerSeq barcode recovery**

Lineage barcode recovery and downstream analysis was performed with custom made bash and python codes. Reads containing a perfect match with tracker-seq barcodes (CTGA..CTG..ACT..GAC..TGA..CTG..ACT..GAC..GACT) were extracted and count tables of tracker-seq barcode occurrence were built for each cell barcode and each UMI. This was done for both the library resulting from the nested PCR strategy, and the gene-expression library (**Extended Data Fig. 7a**). Downstream analysis was performed only on the data obtained from the nested PCR approach. First, we quantified the total number of sequencing reads and the total number of unique UMIs for each cell barcode. Additionally, we computed the mean oversequencing value for each cell barcode, defined as the average number of reads detected for each UMI and each tracker-seq barcode. Then, we selected cell barcodes with more than 100 and 1000 total reads in experiment 1 and 2, respectively (represented by dashed red lines in **Extended Data Fig. 7b**). This criterion automatically selects cells in which at least one tracker-seq barcode was found to be oversequenced with an average maximum of 25 or 10 times (see dashed blue lines in **Extended**

**Data Fig. 7e,f**). Next, for each remaining cell barcodes we calculated the percentage of read counts (normalized to total reads) and observed UMIs (normalized to total observed UMIs) per tracker-seq barcodes, and only kept tracker-seq barcodes that are present at least 10% for both read count and UMI count fractions in at least one cell barcode. For each cell barcode, we extracted the maximum oversequencing value for each tracker-seq barcode and calculated the fraction of counts using only these values. We removed tracker-seq barcodes with a fraction value below 0.2. If more than one tracker-seq barcode still remained per cell, we pooled together tracker-seq barcodes that are less than 4 edits away from each other. If still the case that more than one tracker-seq barcode remained per cell, we assumed that these should be detected according to a multinomial distribution. Therefore, if the fraction of counts for each resulting tracker-seq barcode was lower than 1/N-(1/N)**2 (mean-variance), we deleted it from the pool in that cell. For each cell, the final clonal barcode was defined as the union of the organoid ID from where it was derived and the tracker-seq barcode.

**Consensus non-negative matrix factorization (cNMF) and referencing**

Gene expression programs in tumour scRNA-seq data were inferred using cNMF (v1.4.1)[119] as described in the vignette. In short, cNMF was performed using 100 iterations of NMF with different random seeds for each value of k, the number of components, from 5 to 9. For each value of k, metrics indicating stability, Silhouette score and Frobenius error, were calculated and the k maximizing the Silhouette score and minimizing the Frobenius error was selected for further processing. Outlier components from the selected k were filtered by removing components with a higher mean distance to most similar component of 0.02, which resulted in a program activity matrix and a gene scores matrix. For each malignant cell, program activity values for each component of the selected k were added as metadata to the Seurat object and classified based on highest program activity. Subsequently this annotation was used for further analysis. To compare the meta programs to our previous annotations we used "scclusteval"'s (v.1.0), used for cluster stability and similarity evaluation. The heatmap was generated via "PairWiseJaccardSetsHeatmap" function. Accordingly, fraction of louvian clustering per cNMF programs was compared by using "dittoBarPlot" function using shared nearest neighbour resolution grouped by meta programs.

**Upset plot generation**

Upset plots were used to represent the intersections between the different clonal families and cNMF modules. Only clonal families found in more than one cNMF module are shown. Upset plots were depicted using the function plot from the upsetplot python package.

**Comparison of large versus small clonal families**

The fraction of each clonal family was computed for each sample. We defined as big clones the clones that are detected in more than 20% of all the cells from each sample. Sample DMG0155 was removed from this analysis as it presented as only one large clone (>99% of the cells had the same tracker-seq barcode). We then perform DEG analysis comparing the large versus small clones using the function rank_genes_groups from the scanpy python package and selected DEGs (log2fold change >0.7) to perform METASCAPE analysis (selected GO terms included in **Supplementary Table S4c,e**).

#### GD2 CAR T cell expansion and selection

CD8 GD2 CAR T cells (14G2a GD2-4-1BBz CAR) and donor-matched mock-transduced CD8 T cells, were produced as previously described[120]. CAR T cells and mock transduced T cells were expanded using a rapid expansion protocol[121]. T cells were cultured in RPMI 1640 + GlutaMax (Thermo Fisher, Cat. #61870036), supplemented with 2.5-10% human serum (Sanquin), 1% Pen-Strep, and 0.5M beta-2-mercaptoethanol (Thermo Fisher, Cat. #21985023), on a feeder cell mixture comprising of sub-lethally irradiated allogenic PBMCs, Daudi, and LCL-TM cells, in the presence of 50 U/ml IL-2 (R&D Systems, Cat. #P60568), 5 ng/ml IL-15 (R&D Systems, Cat. #P40933), and 1 µl/ml PHA-L (Sigma-Aldrich, Cat. #11249738001) and cryopreserved after 14 days of expansion. Prior to experiments, T cells were thawed and rested in RPMI 1640 + GlutaMax, with 10% Fetal Bovine Serum (FBS, Thermo Fisher, Cat. #10500064) and 1% Pen-Strep, supplemented with 50 U/ml IL-2 (Miltenyi, Cat. #130-097-743), 2000 U/ml IL-7 (Miltenyi, Cat. #130-095-367), and 50 U/ml IL-15 (Miltenyi, Cat. #130-095-760) for 3 days at 37°C with 5% $CO_2$. For selection of GD2 CAR T cells based on NCAM1 expression, a similar expansion protocol was used, but without addition of IL-15 and Daudi cells. After resting, cells were washed and stained for 30 min at 4˚C in FC buffer with LIVE/DEAD Fixable Near-IR Dead Cell Stain (1:1000; Thermo Fisher), CD3-APC (1:80; BD BioLegend, clone SK7) and NCAM-1-HiLyte-488 (1:200; QVQ, FSH-10B10). $CD3^{+}$ $NCAM1^{-}$ or $CD3^{+}$ $NCAM1^{+}$ GD2 CAR T cell populations were separated by FACS on a BD FACSAria II Cell Sorter and immediately used for experiments.

#### GD2 CAR T cell treatment

Four months after tumour induction, DMGOs were transferred to 12-well suspension plates and untreated or treated with 500,000 $CD8^+$ GD2 CAR T cells, or mock transduced $CD8^+$ T cells per DMGO. 7 days after the start of treatment, 500,000 T cells were added per DMGO for a second round of treatment. Tumour size during treatment was monitored by imaging on day 0, 3, 7, 10, and 14, or on day 0, 2, 5, 7, 10 and 14 for treatment with NCAM1 selected T cells using a Leica Thunder DMi8 microscope with a 10X objective. In addition, one DMGO was treated on day 0, 8 and 15 with GD2 CAR T cells and imaged on day 8, 15, 28 and 35. After THUNDER software-mediated computational clearing of the imaging data, tumour size for each time point was quantified using Fiji. In short, background signal, defined as GFP-negative areas within the organoid, was subtracted. The organoid surface was set as region-of-interest (ROI) and mean gray values of the GFP channel for the ROI were calculated. Additionally, at day 7, 10 and 14 supernatant of the co-cultures was collected and IFNγ concentration was measured with ELISA (R&D Systems, Cat. #DY285B).

Similarly, untransformed BrOs were transferred to 12-well suspension plates and treated with 500,000 $CD8^+$ GD2 CAR T cells per BrO. 7 days after the start of treatment, 500,000 T cells were added per BrO for a second round of treatment. Organoid appearance during treatment was monitored by imaging on day 0 and 14.

##### FACS of GD2 CAR T cells and DMGO GD2 expression

DMGOs treated with GD2 CAR T cells were dissociated 14 or 35 days after initial T cell addition with the Neural Tissue Dissociation Kit (P) (Miltenyi Biotec, Cat. #130-092-628), as described above for preparation of single cell RNA and tracker seq libraries. Dissociated cells were washed and stained in FC buffer with CD3-APC (1:80; BD Biosciences, clone SK7) and LIVE/DEAD Fixable Near-IR Dead Cell Stain (1:1000; Thermo Fisher) for 30 min at 4˚C. $CD3^+$ T cells and $GFP^+$ tumour cells were sorted on a CytoFLEX SRT Benchtop Cell Sorter (Beckman Coulter) and immediately processed for scRNA-seq. To confirm DMGO GD2 expression for GD2 CAR T cell treatment evaluation, a day 60 post-electroporation DMGO sample was dissected for the tumour region to enrich for tumour material, mechanically dissociated, and cultured for 2 additional weeks to expand tumour cells. Cells were retrieved from the culture plate using StemPro Accutase (Gibco, Cat. #A1110501) and passed through a 70 µm Flowmi cell strainer (Merck, Cat. #BAH136800070) to create a single cell suspension. Dissociated cells were centrifuged at 500 RCF for 5 minutes at 4 °C and resuspended and washed in FC buffer (2% fetal bovine serum (FBS), 1x PBS). Cells were either left unstained or stained with LIVE/DEAD Fixable Near-IR Dead Cell Stain (1:1000; Thermo Fisher) and GD2-PE (1:200, clone 14.G2a, BD Biosciences, Cat. #562100) for 30 min at 4˚C. After staining, cells were washed twice in FC buffer, acquired on a Sony SH800s (Sony Biotechnology), and analysed using FlowJo Software (v10.9.0).

**Pre-processing and analysis of GD2 CAR-T cell scRNA seq datasets**

As the first quality control step, doublets (two, or more, cells captured in the same droplet) for each sample were identified and removed using the scDblFinder package[122], with default settings. Low quality cells with high mitochondrial content (> 15%), or cells with extremely high or low reads (< 200 genes or > 6500 genes), or cells with extremely high reads (> 35000 reads) were removed. Normal Seurat V4[123] workflow was used to normalize and scale reads, and the 3000 most variable features determined using "FindVariableFeatures" in the Seurat package. Cell cycle confounding effect was eliminated from the dataset via the removal of cell cycle-related genes from the variable features of the dataset. PCA was performed using “RunPCA” function. First 30 PCs were used for non-linear dimensionality reduction utilizing UMAP[124] method, implemented via “RunUMAP” function of the Seurat package. Clustering analysis was performed on the first 10 PCs using the Seurat package's `FindNeighbors` and `FindClusters` functions. A resolution parameter of 0.45 was applied, and the original Louvain algorithm was used. To identify sub-populations, marker genes for each cluster were determined through the `FindAllMarkers` function. Markers obtained from this analysis were then examined to profile genes associated with known CD8 T cell subsets, as well as to project previously published signatures (see T cell signature projection below). Only markers with adjusted p values below 0.05 were taken into consideration. In addition, DEGs were used as input for gene ontology (GO) enrichment analysis using the GO resource (https://geneontology.org).

To integrate three different NCAM1-sorted, unexposed, and bulk RNA sequencing experimental batches, we applied the Seurat-based Canonical Correlation Analysis (CCA) integration method. As integration features, we used 1000 variable features from each dataset, along with differentially expressed genes between conditions (min.pct =0.3, log fold change =1.5), to account for biological variability. To assign cell populations to the clusters identified in **Fig. 3e**, we estimated the proportion of cells from each cluster described in that figure for every cluster of the integrated dataset. Newly emerging populations were defined based on differentially expressed markers and their origin, categorized by whether they originated from exposed, unexposed, $NCAM1^+$ or $NCAM1^-$ populations.

To determine the identity of the $T_{MA}$ cluster, we mapped our GD2 CAR T cell subsets to the $CD8^+$ TIL subsets from the Chu *et al*. resource dataset[5] using Seurat’s FindTransferAnchors(). The GD2 CAR T cell identities were then transferred to the Chu *et al*. dataset with TransferData(), retaining only high-confidence predictions (score > 0.5). These transferred identities were used to calculate the proportion of $T_{MA}$ GD2 CAR T cells within each $CD8^+$ TIL subset from the Chu *et al*. dataset.

**T cell signature projection**

To evaluate the expression of established T cell signatures in GD2 CAR T cell scRNA seq datasets, we used a gene signature specific to serial killer engineered T cells that we previously obtained (Dekkers *et al*.[67], Supplementary Table 4). Utilizing the VISION R package[125], we computed and visualized the overall enrichment of the identified gene set atop UMAP cell embeddings of our dataset. In addition, we projected our own GD2 CAR T cell signature profiles onto a pan-cancer CD8 tumour infiltrating lymphocyte (TIL) atlas from Chu *et al.*[5], which encompasses T cells infiltrating brain tumours. For each GD2 CAR T cell subset, markers obtained through DEG analysis were meticulously curated to obtain the most relevant markers, ensuring an adjusted p-value below 0.00001. Accessing a publicly available and interactive online data portal (https://singlecell.mdanderson.org/TCM/), we acquired a rds file containing the Seurat object pertinent to scRNA seq data of CD8 TILs. Subsequently, the VISION package was employed to perform the projection of our GD2 CAR T cell signatures onto this dataset.

**Primitive macrophage progenitor generation and integration**

The protocol to generate primitive macrophage progenitors (PMPs) was adjusted from Gutbier *et al.* (2020). In short, 70-80% confluent H1 stem cells were detached with Gentle Cell Dissociation Reagent (GCDR, #100-0485, StemCell). For EB formation, 7000 cells were plated per well of an ultra-low attachment treated U-bottom 96-well plate (#650970, Greiner Bio-One) in mTeSR+

(#100-0276, StemCell) medium, containing 50 uM ROCK inhibitor (Y27632, #M1817, AbMole), 50 ng/ml BMP4 (#78211, StemCell), 50 ng/ml VEGF (#100-20-100ug, PeptroTech,) and 20 ng/ml SCF (#130-093-991, Miltenyi Biotec). On day 2, fresh medium was added to each well. On day 4, EBs were transferred to a 6-well plate with X-VIVO 15 (#BE02-060F, Lonza) medium, containing 1X GlutaMax (Gibco), 1X Pen-Strep (Gibco), 100 ng/ml M-CSF (#300-25-50ug, PeptroTech), and 25 ng/ml IL-3 (#AF-200-03-10ug, PeproTech). The medium was refreshed once a week. After about 3 weeks, the release of PMPs in the supernatant was observed. PMPs were collected from the supernatant and counted, before adding 100-200k cells per brain organoid in maturation medium (1:1 Advanced DMEM/F-12 (Gibco) and Neurobasal (Thermo Fisher) medium, 1X GlutaMax, 0.5x N-2 (Gibco), 0.5X B27 without vitamin A (Gibco), and 1X Pen-Strep). Brain organoids with PMPs were kept on a microtiter orbital shaker inside a 37°C/5% CO2 incubator for 1-3 weeks for integration and differentiation of the PMPs to microglia. Alternatively, brain organoids were sectioned into 200 μm thick slices using a vibratome, transferred to a 24-well suspension plate in 750 μL maturation medium and incubated at 5% CO2 and 37˚C for 3 days. Then, 50-200k PMPs were added per slice in 750 μL maturation medium in a 24-well suspension plate for 7 days before starting treatment. For GD2 CAR-T cell treatment, 200k $CD8^+$ GD2 CAR T cells were added in 750 μL maturation medium per 24-well to the slices. Tumour size during treatment was monitored by imaging on day 0, 3, 7, 10, and 14, using a Leica DMIL LED FLUO microscope with a 10X objective.

**SORT-seq of microglia and GD2 CAR T cells**

BrOs and DMGOs containing microglia and optionally treated with GD2 CAR T cells were dissociated 21 days after initial microglia incorporation with the Neural Tissue Dissociation Kit

(P) (Miltenyi Biotec, Cat. #130-092-628), as described above for preparation of single cell RNA and tracker seq libraries. Dissociated cells, control PMPs and unexposed GD2 CAR T cells were washed and stained in FC buffer with CD3-BV421 (1:100; BD Biosciences, clone SK7) and LIVE/DEAD Fixable Near-IR Dead Cell Stain (1:1000; Thermo Fisher) for 30 min at 4˚C. $CD3^+$ T cells and $mScarlet^+$ microglia/ PMP were sorted into 386-well plates containing well-specific barcoded primers (Single Cell Discoveries), one cell per well, on a Sony SH800s (Sony Biotechnology). Plates containing sorted cells were immediately snap frozen on dry ice and processed for SORT-seq by Single Cell Discoveries. In short, scRNA-seq was performed according to an adapted version of the SORT-seq protocol[126] with primers described in van den Brink *et al.*[127]. Cells were heat-lysed at 65°C followed by cDNA synthesis. All the barcoded material from one plate was pooled into one library and amplified using *in vitro* transcription (IVT). Following amplification, library preparation was done following the CEL-Seq2 protocol[128] to prepare a cDNA library for sequencing using TruSeq small RNA primers (Illumina). The DNA library was paired-end sequenced on an Illumina Nextseq™ 500, high output, with a 1×75 bp Illumina kit (read 1: 26 cycles, index read: 6 cycles, read 2: 60 cycles).

**Microglia phagocytosis assay**

To assess the functionality of the integrated microglia in the brain organoids, a myelin phagocytosis assay was performed. Brain stem organoids were sliced and mScarlet-labeled PMPs were added as described above. CSFE-labeled myelin debris, kindly provided by the Akkari lab[82], was injected into the slice using a glass needle and a FemtoJet 4i. The slices were immediately imaged on a Leica STELLARIS microscope at 37oC and 5% CO2 overnight with a time interval of 5-10 minutes.

**Luminex analysis of the culture supernatant**

Determination of protein concentrations was done by Luminex, as previously published[129]. In short, acquisition of data was performed using a FLEXMAP 3D system (Bio-Rad) using xPONENT 4.3u1 software (Luminex). Data analysis was performed using Bio-Plex Manager 6.2 (Bio-Rad). All assays were performed at the ISO9001:2008 certified MultiPlex Core Facility of the University Medical Center Utrecht.

**Statistical and heatmap analysis**

Statistics on bulk sequencing data was computed by built-in functions of R ("stats", v4.3.1) using one-way ANOVA with post-hoc Tukey Honest significance difference. PCA, Spearman's Rank and gene expressions were plotted using ggplot2(v3.4.2), heatmaps were generated using pheatmap package (v1.0.12). Statistics on electroporation efficiency and tumour induction was calculated using the two-tailed independent t-test (function: t.test). For each batch and timepoint the mean and standard deviation was calculated, individual values were imported and plotted in GraphPad Prism (v.8.0.2) using summarizing stacked bar plots. All statistical tests have been performed with the assumption of a normal distribution, equal variance per sample and a confidence interval of at least 95% (alpha = 0.05). To evaluate tumour response to GD2 CAR T cells in the presence or absence of microglia (**Fig. 4k**), the normalized tumour area at each timepoint was analysed using a linear mixed-effects model, accounting for fixed and random effects related to batch and organoid variation. Multiple models were tested, and the best-fitting model was selected.

#### Data availability

All used R and Python scripts are available in our laboratory GitHub. All sequencing datasets (bulk and single cell) will be deposited on NCBI Gene Expression Omnibus (GEO) before publication. Sequencing metadata is provided in **Supplementary Table S2**.

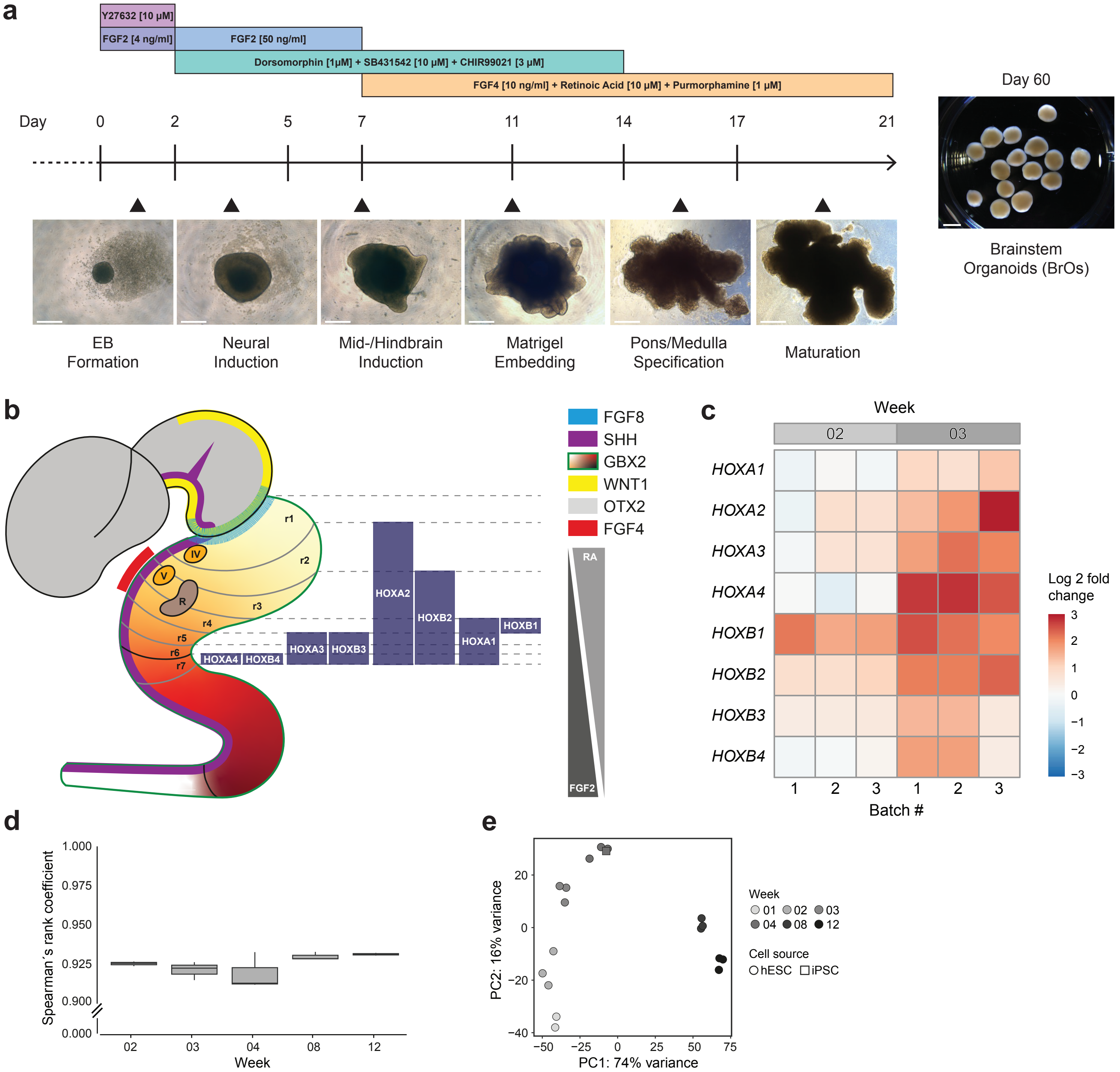


**Extended Data Figure 1**

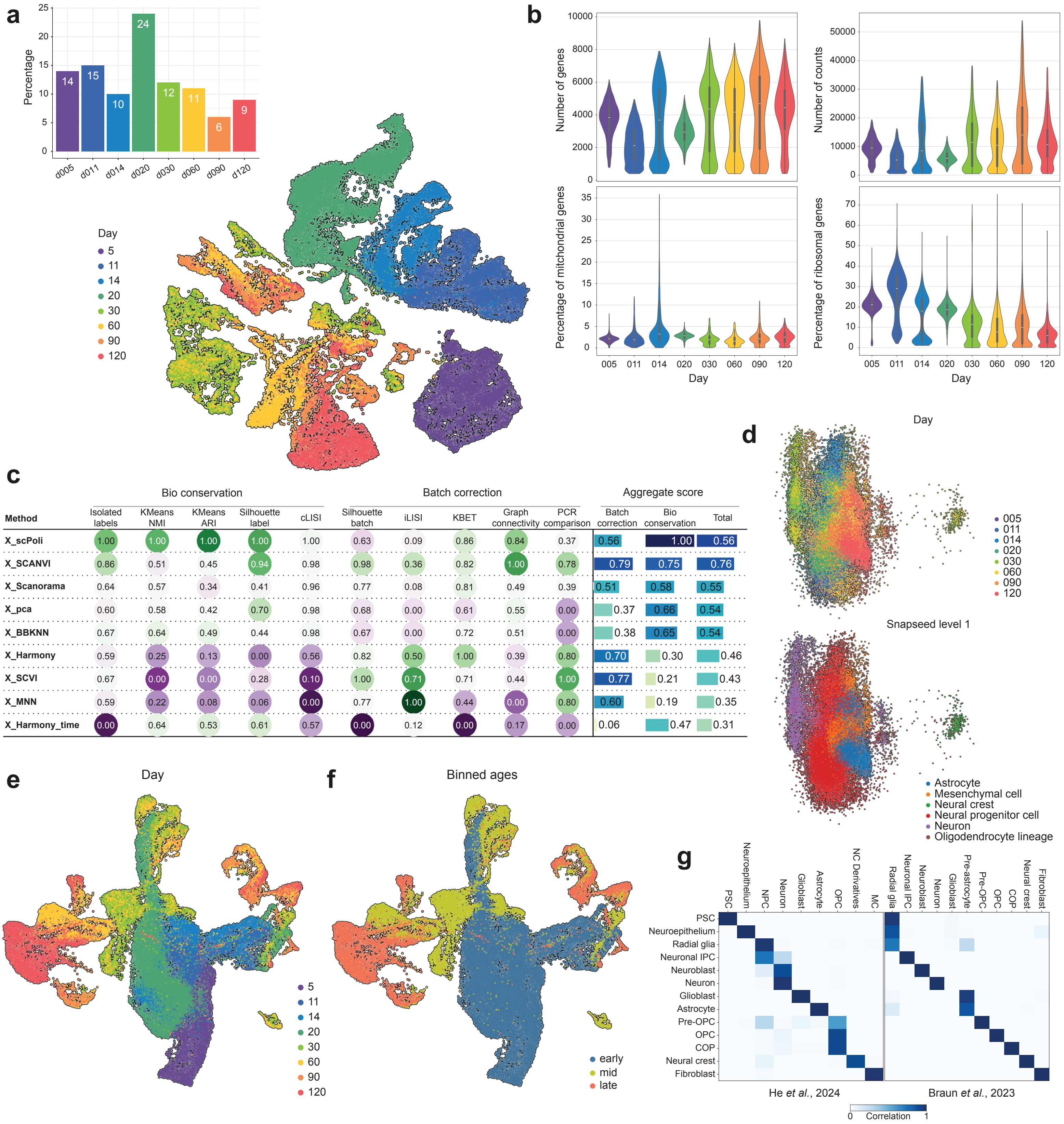


**Extended Data Figure 2**

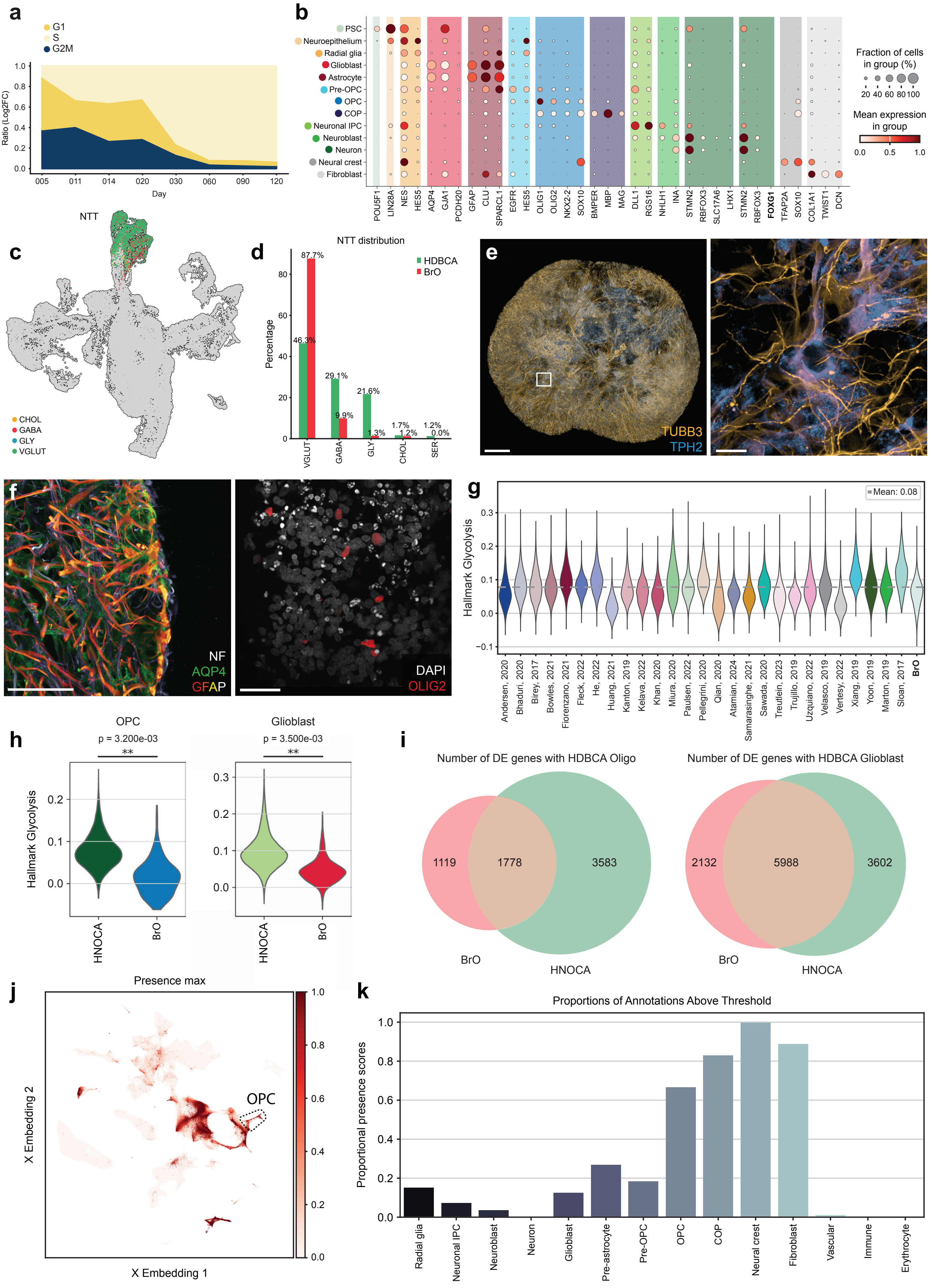


**Extended Data Figure 3**

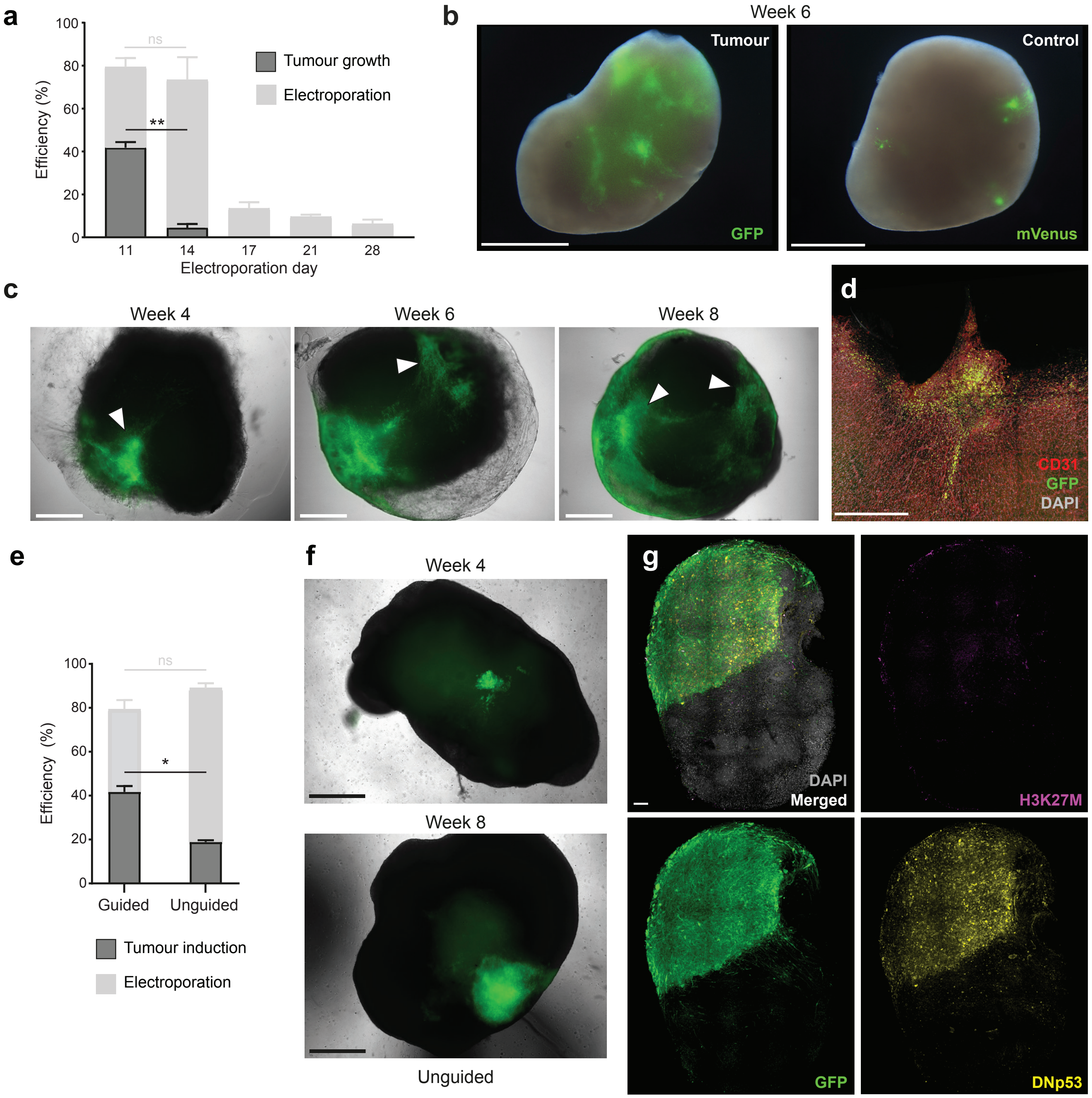


**Extended Data Figure 4**

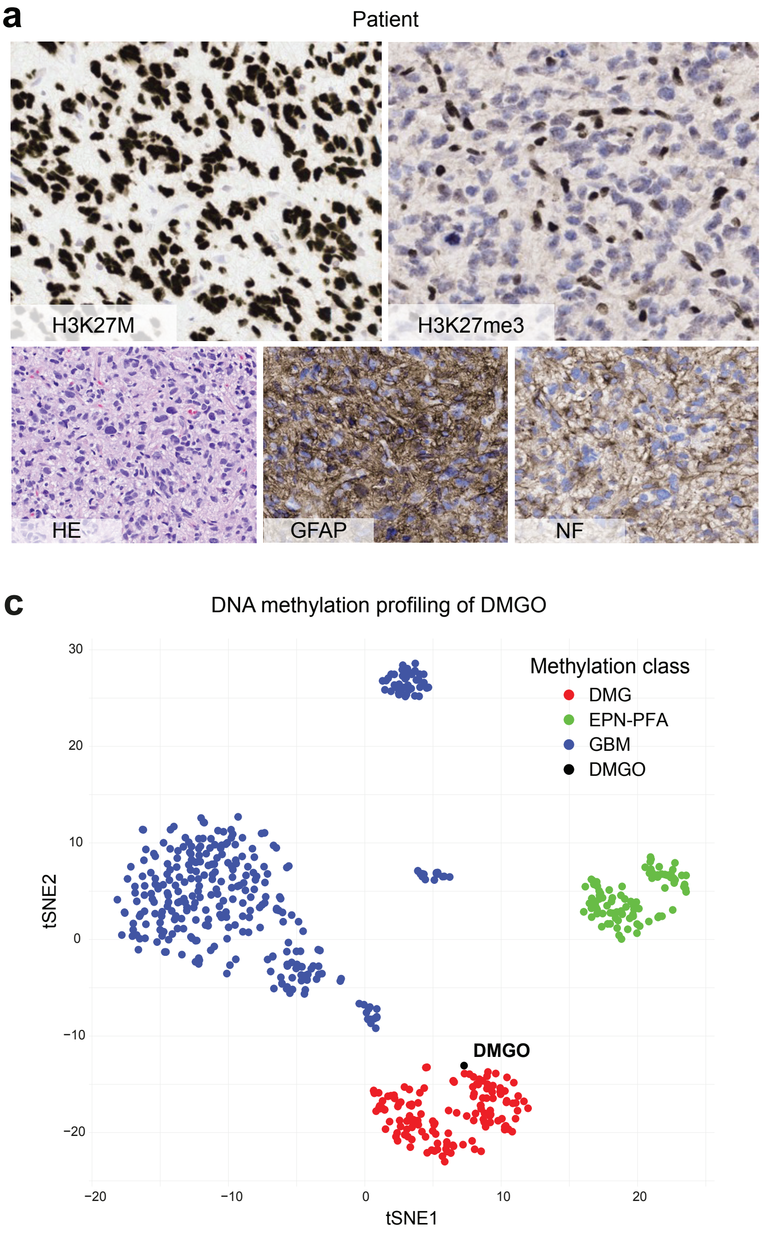


**Extended Data Figure 5**

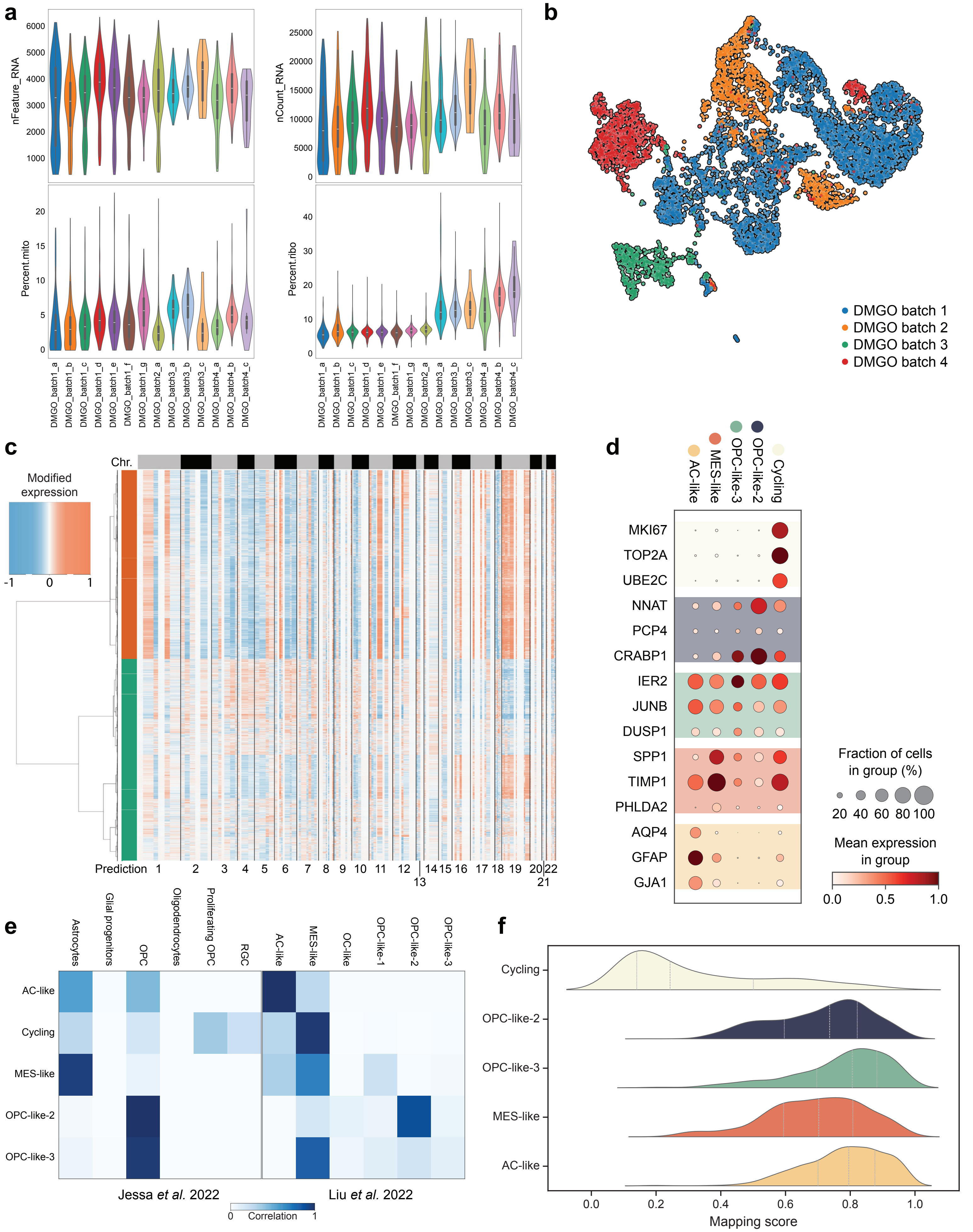


**Extended Data Fig. 6**

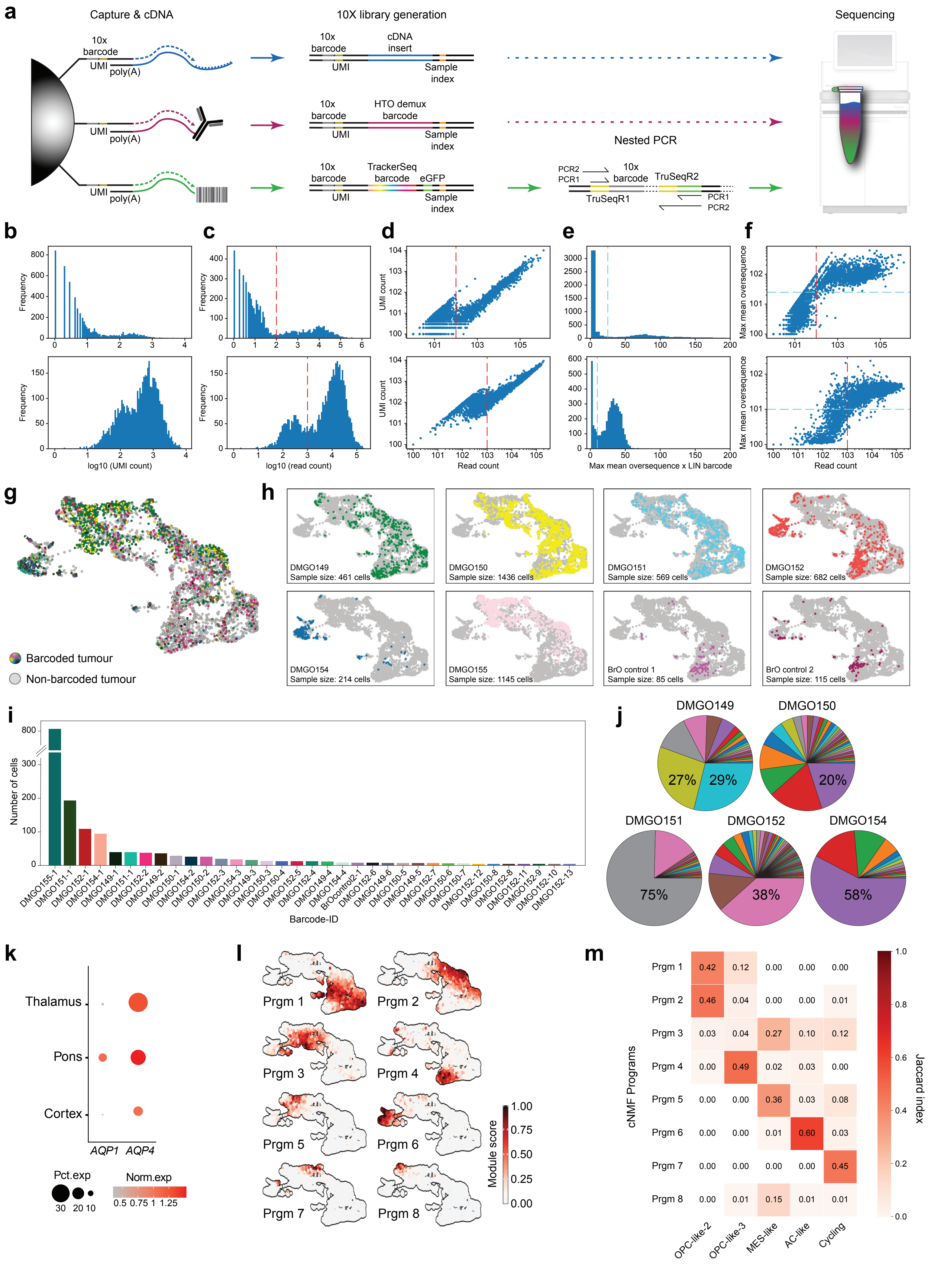


### Extended Data Figure 7

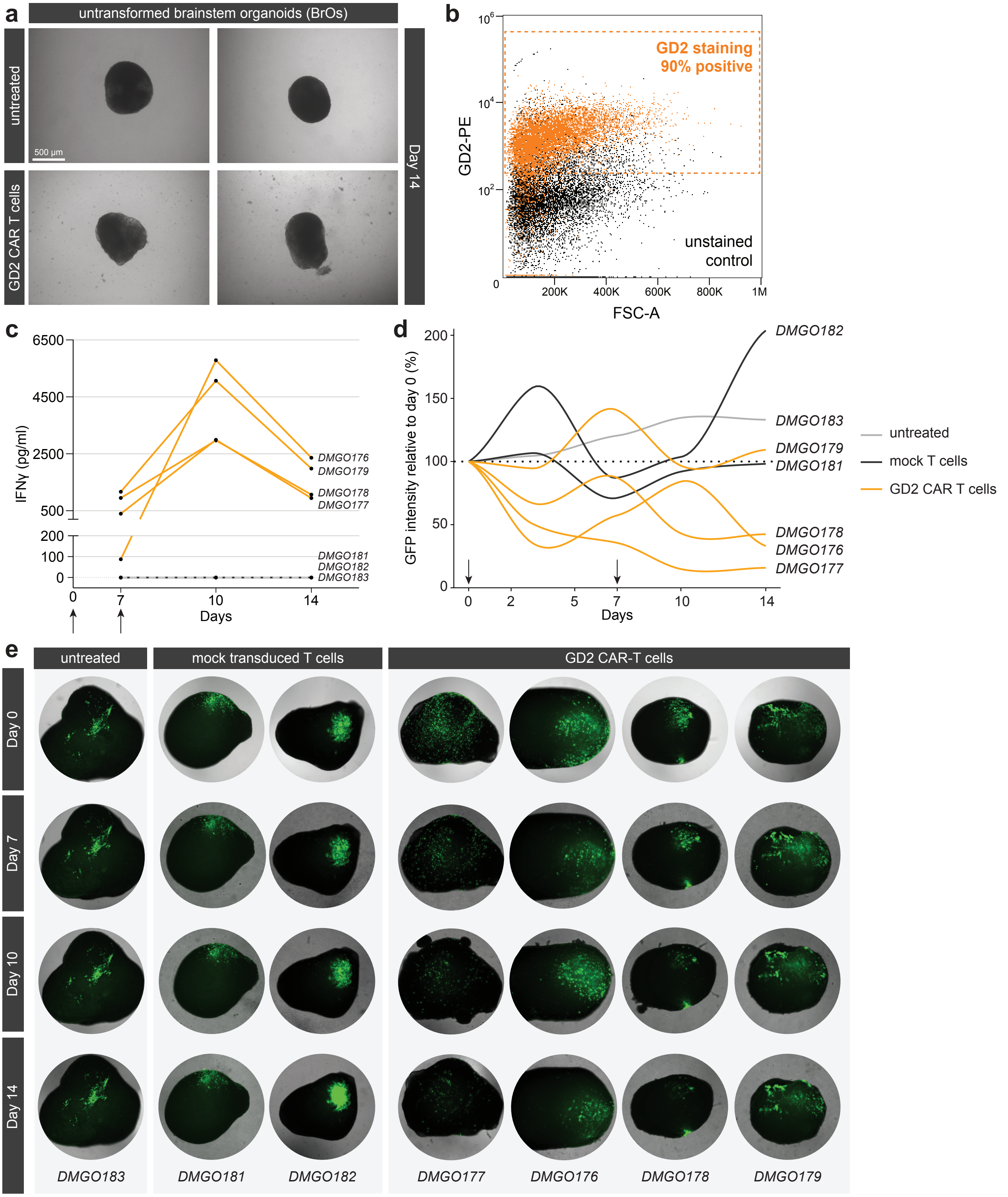


**Extended Data Figure 8**

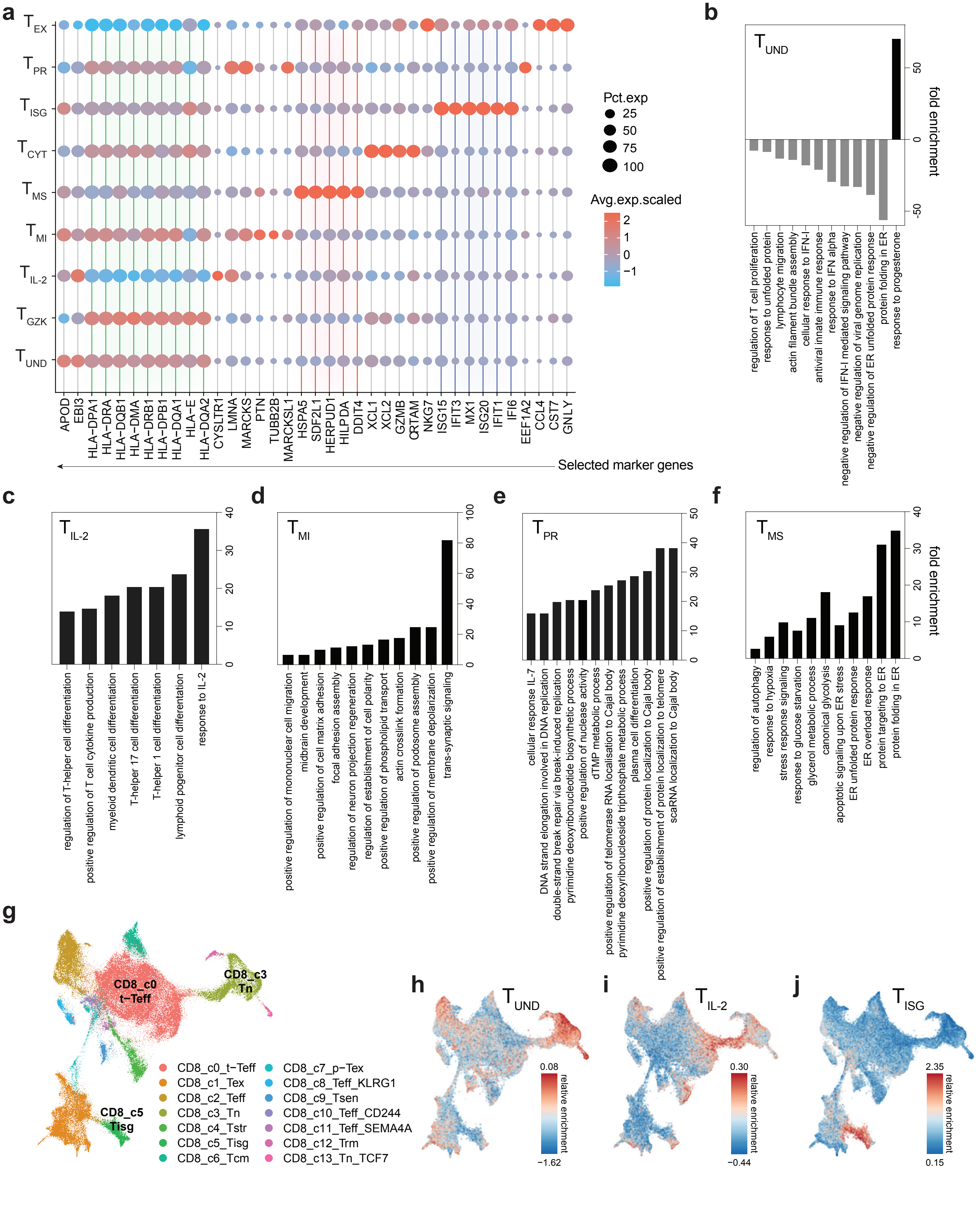


**Extended Data Figure 9**

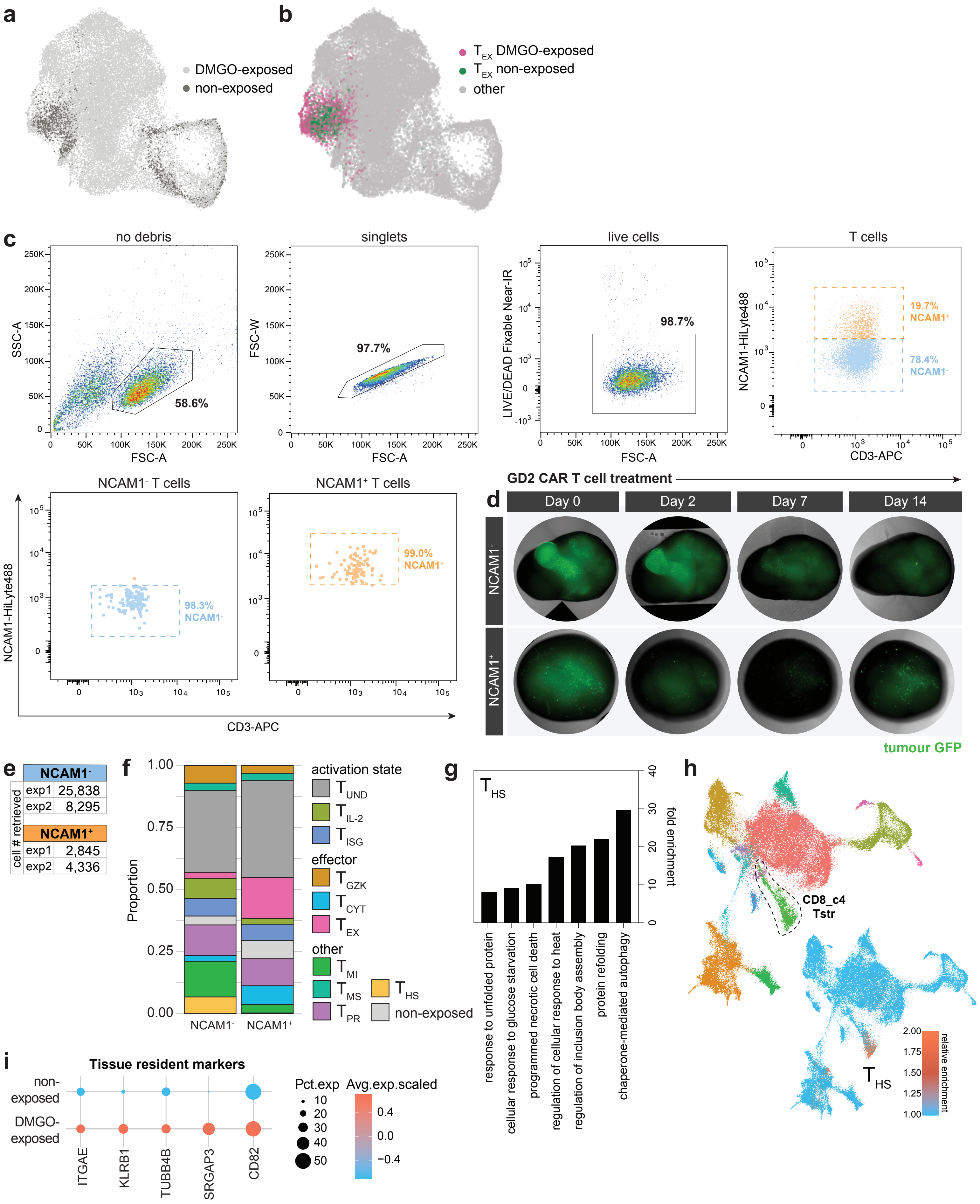


**Extended Data Figure 10**

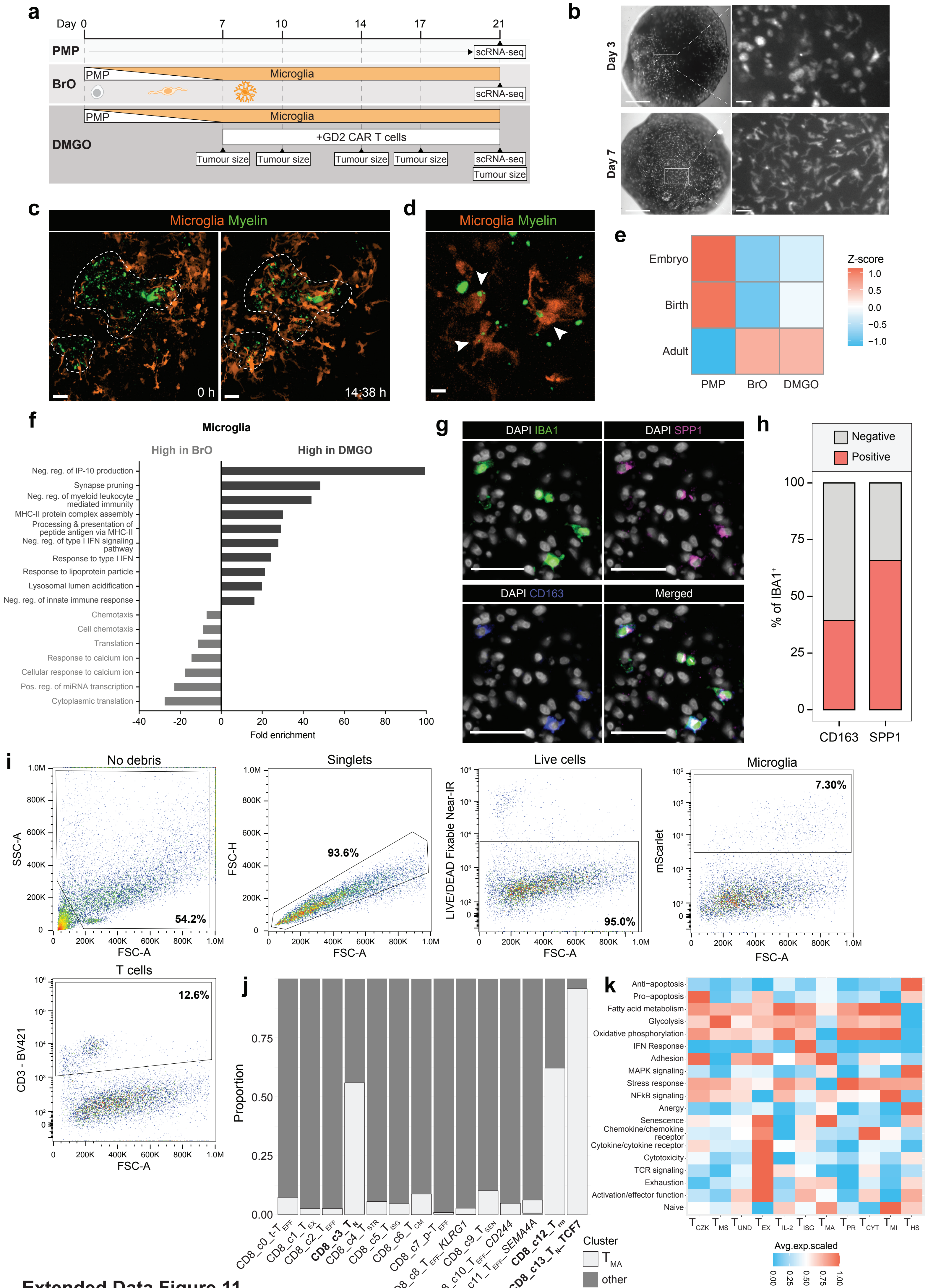


**Extended Data Figure 11**
